# Parallel functional testing identifies enhancers active in early postnatal mouse brain

**DOI:** 10.1101/2021.01.15.426772

**Authors:** Jason T. Lambert, Linda Su-Feher, Karol Cichewicz, Tracy L. Warren, Iva Zdilar, Yurong Wang, Kenneth J. Lim, Jessica Haigh, Sarah J. Morse, Cesar P. Canales, Tyler W. Stradleigh, Erika Castillo, Viktoria Haghani, Spencer Moss, Hannah Parolini, Diana Quintero, Diwash Shrestha, Daniel Vogt, Leah C. Byrne, Alex S. Nord

**Author notes:** These authors contributed equally to this work (Jason T. Lambert, Linda Su-Feher). Correspondence to Alex S. Nord.

## Abstract

Enhancers are cis-regulatory elements that play critical regulatory roles in modulating developmental transcription programs and driving cell-type specific and context-dependent gene expression in the brain. The development of massively parallel reporter assays (MPRAs) has enabled high-throughput functional screening of candidate DNA sequences for enhancer activity. Tissue-specific screening of *in vivo* enhancer function at scale has the potential to greatly expand our understanding of the role of non-coding sequences in development, evolution, and disease. Here, we adapted a self-transcribing regulatory element MPRA strategy for delivery to early postnatal mouse brain via recombinant adeno-associated virus (rAAV). We identified and validated putative enhancers capable of driving reporter gene expression in mouse forebrain, including regulatory elements within an intronic *CACNA1C* linkage disequilibrium block associated with risk in neuropsychiatric disorder genetic studies. Paired screening and single enhancer *in vivo* functional testing, as we show here, represents a powerful approach towards characterizing regulatory activity of enhancers and understanding how enhancer sequences organize gene expression in normal and pathogenic brain development.

## INTRODUCTION

*Cis*-regulatory elements such as enhancers are critical drivers of spatiotemporal gene expression within the developing and mature brain^1^. Enhancers integrate the combinatorial functions of transcription factors^2^ and chromatin organizers^3^ to drive cell and regional-specific expression of genes across brain development. DNA sequence variation within enhancers has been associated with both evolution^4^ and the genetic etiology of neurological disorders^5^. The first identified enhancers were defined by functional capacity to amplify transcriptional activity in reporter plasmids^6, 7^. Putative enhancers have been predicted via DNA conservation using comparative genomics and, more recently, by epigenetic signatures such as open chromatin from DNaseI hypersensitive site sequencing (DNase-seq) or assaying for transposase-accessible chromatin using sequencing (ATAC-seq)^5^, histone tail modifications from chromatin immunoprecipitation sequencing (ChIP-seq)^1, 8^, and 3D chromatin organization^9^; however, these approaches are proxy measurements that do not directly evaluate whether a DNA sequence acts as a functional enhancer^10^.

Enhancer reporter assays assess the ability of candidate DNA sequences to drive expression of a reporter gene, and have been used as the primary means of functionally testing activity of predicted enhancers^11^. Transgenic mouse enhancer assays have been applied to characterize the regulatory activity of candidate DNA sequences in the mouse, in both the developing and mature brain^1, 8, 12^. These assays offer exceptional information on tissue-specific enhancer activity but require one-by-one testing of individual candidates. The advancement of massively parallel reporter assays (MPRAs) has led to the ability to functionally screen a multitude of DNA sequences for enhancer activity in parallel within a single experiment using sequencing-based quantification^13^. To date, there are a few promising demonstrations of MPRA applied *in vivo* in brain^14, 15^, though screening approaches to characterize enhancers remain highly novel, particularly in developing brain. Functional assessment of enhancer activity and the use of enhancers to express reporters and genes in mouse brain cells has emerged as an area of major interest^16–19^, and screening approaches represent an ideal approach to perform broad testing at scale.

Here, we adapted a functional enhancer screening approach, self-transcribing active regulatory region sequencing (STARR-seq)^20^, for application to early postnatal mouse brain via recombinant adeno-associated virus (rAAV) delivery. We screened a library of candidate human DNA sequences spanning putative brain-specific regulatory sequences and regions associated with genome-wide association studies of epilepsy and schizophrenia. We identified sequences capable of driving reporter expression in early postnatal mouse cortex, and validated positive and negative MPRA predictions, including verifying enhancers within the neuropsychiatric disorder-associated third intron of *CACNA1C*^21^. Our results demonstrate the utility of parallel functional testing to dissect the regulatory structure of enhancers in early postnatal mouse brain and highlight opportunities for enhancer screening in studies of normal brain development and function as well as the etiology of neurodevelopmental and neuropsychiatric disorders.

## RESULTS

### Cloning and AAV packaging of MPRA library

We generated an MPRA library targeting 408 candidate human DNA sequences for testing, each ∼900 bp in size, to assess functional enhancer activity *in vivo* in early postnatal mouse brain. These sequences were categorized into four groups, two groups without ascertainment based on potential regulatory activity, and two groups selected based on presence of epigenomic signatures commonly used to predict enhancer activity (Supplementary Table 1). For the first group (“GWAS”), we selected genomic intervals containing single-nucleotide polymorphisms (SNPs) identified as lead SNPs from genome-wide association studies (GWAS) for epilepsy^22^ and schizophrenia^23^ as identified using the NHGRI-EBI GWAS Catalog^24^. For the second group (“LD”), we selected five lead SNPs from these same studies, and included genomic intervals that included all SNPs in linkage disequilibrium (LD, r^2^ > 0.8, 1000 Genomes Project^25^) within these non-coding regions associated with the *CACNA1C* and *SCN1A* loci. For the third group (“FBDHS”), we selected regions overlapping DNaseI hypersensitive sites in fetal human brain from ENCODE^5^ within copy number variant regions near autism risk genes^26, 27^. Finally, we included human sequences (“PutEnh”) that were predicted to have either enhancer activity or no activity in postnatal mouse brain based on previous transgenic mouse enhancer assays and on mouse brain H3K27ac ChIP-seq data^1, 8^.

We used a modified STARR-seq^20^ MPRA orientation, in which the candidates of interest were cloned in the 3’ untranslated region (3’ UTR) of a reporter gene (here EGFP) driven by the minimal promoter, Hsp68^28^ (Figure 1A). We cloned all regions of interest (referred to from here on as amplicons) from pooled human DNA via polymerase chain reaction (PCR) and Gibson assembly directly into a viral DNA plasmid backbone containing the enhancer reporter and inverted terminal repeat regions (ITRs) necessary for self-complementary adeno-associated viral (scAAV) packaging (Supplemental Figure 1). Following batch cloning, we sequenced the pooled plasmid library, verifying presence of 345 candidate amplicons, and we packaged this into an scAAV library using AAV9(2YF)^29^. AAV9(2YF) is a tyrosine-mutated derivative of AAV9, which has increased transduction in the brain and similar tropism compared to standard AAV9. scAAV has the advantage of faster transduction rates compared to conventional AAV^30^.

**Figure 1.**
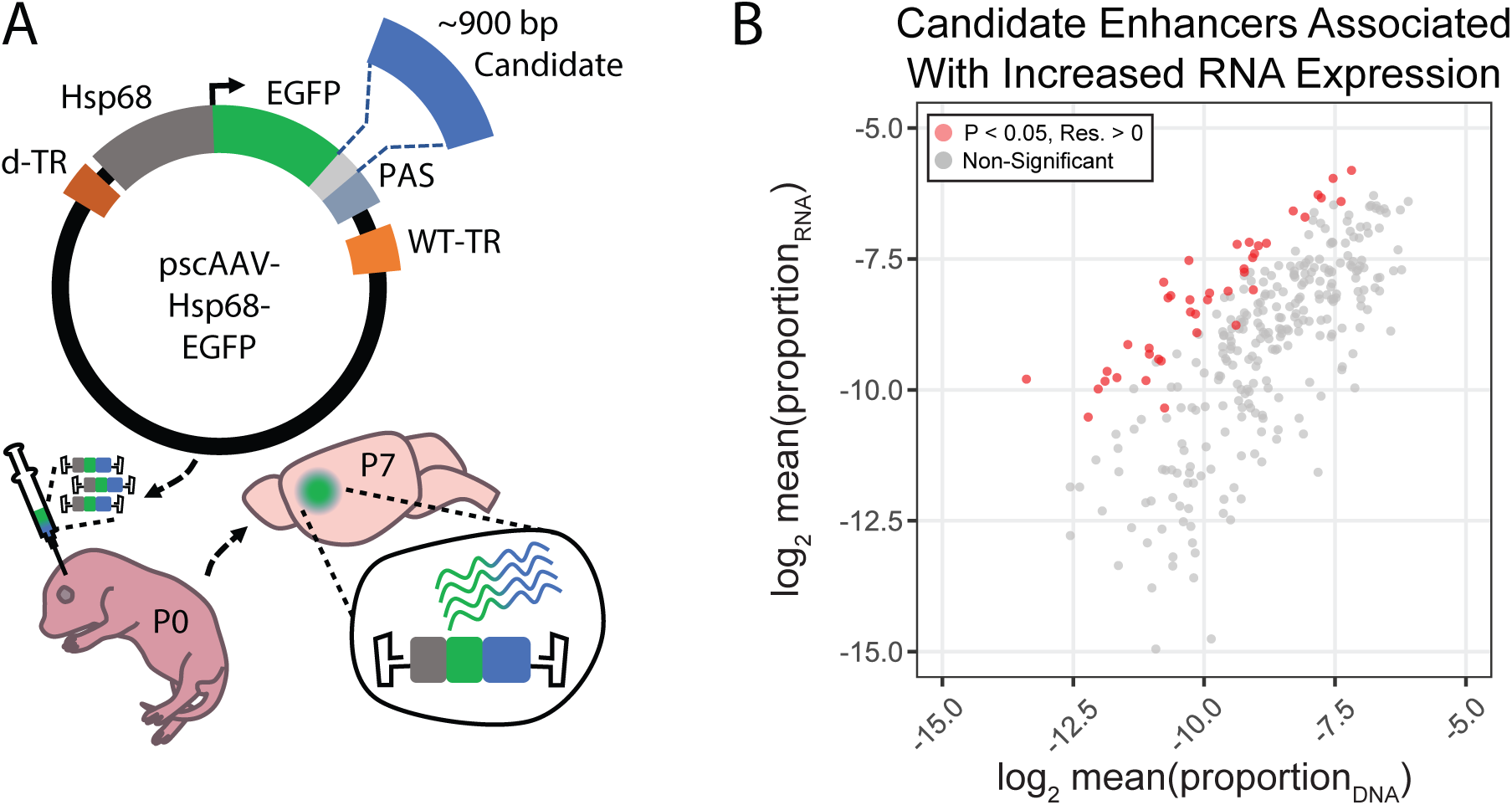
In vivo parallelized functional enhancer reporter screen. (**A**) Schematic of in vivo parallelized functional enhancer reporter assay. The test library was generated using the previral vector pscAAV-Hsp68-EGFP, which contained a multiple cloning site (light grey) between the EGFP reporter and polyadenylation site (PAS). Purified PCR amplicons of test amplicons were cloned into the vector using Gibson assembly. The previral library was packaged into AAV9(2YF), and the viral library delivered to the brain via ventricular injection at postnatal day (P)0. Brains were collected at P7. (**B**) Correlation of mean RNA/DNA ratio in the assay showing amplicons tested by the multiple linear model (N = 308). Amplicons with significantly (P < 0.05) increased model residual value (Res.) in RNA compared to DNA are shown in red.

### Application of *in vivo* MPRA and assessment of reproducibility of biological replicates

We delivered the viral library to the left hemisphere of four neonatal mouse brains at postnatal day (P)0 via intraventricular injection and collected forebrain tissue seven days after transduction at P7 (Figure 1A). We isolated viral genomic DNA, representing input or delivery control, as well as total RNA and generated amplicon sequencing libraries for the left hemispheres for each replicate as well as for the previral batch-cloned plasmid library. Following sequencing, de-duplicated aligned reads to the human genome were used to generate amplicon summary counts (Supplementary Table 2). We observed significant correlation between biological replicates (Supplemental Figure 2A, B). We also observed strong correlation between amplicon read counts in the pooled previral plasmid library (“Library”) and genomic AAV DNA (“DNA”) collected from each injected P7 brain (Supplemental Figure 2A), with 308/345 (89%) passing count and amplicon library proportion quality control thresholds (see Methods). To test the effect of PCR bias during library preparation on amplicon counts, we generated one technical RNA replicate, Sample 4-35, using a higher cycle count (Supplemental Figure 2B), and found similar amplicon representation and dropout patterns as its matched lower-cycle technical replicate. Taken together, this demonstrates that viral packaging, neonatal delivery, P7 sample collection and processing, library generation, and sequencing did not substantially affect amplicon representation. Although correlation between amplicons generated from P7 cDNA (“RNA”) was strong overall, particularly for amplicons with robust cDNA expression, each replicate included a small subset of lower cDNA expression amplicons that showed replicate-specific cDNA dropout (Supplemental Figure 2B). Amplicon representation in the previral library and P7 viral DNA was correlated with amplicon GC content (Supplemental Figure 3A-B), indicating GC-based differences impact PCR and cloning efficiency. There was reduced strength in correlation between GC content and the MPRA RNA (Supplemental Figure 3C-D) and no further GC bias arose between viral packaging, delivery, and recapture of the library, suggesting that GC bias originated from the batch cloning process. Input viral DNA and MPRA RNA amplicon counts were also generally correlated, indicating basal MPRA transcription independent of specific enhancers, as reported in other MPRA studies^31, 32^.

### Identification of putative enhancer activity in P7 brain

First, we performed quality control and filtering, removing amplicons with DNA and RNA representation that was too low for statistical analysis (N = 308, Supplementary Table 3). Having confirmed reproducibility of input viral DNA and MPRA RNA collection, we performed activity estimation. We used the middle 80% of amplicons (N = 248) ranked by RNA/DNA ratio (Supplemental Figure 3E) to build a linear model to account for background MPRA RNA based on amplicon representation and GC content (Supplemental Figure 3E-G). For regression-based estimation of activity, amplicon representation across each replicate was combined to generate a summary activity value.

To identify amplicons with activity above expected based on background (i.e. enhancers), we applied the model to the full set of amplicons (N = 308) to generate residual values that represent observed RNA levels compared to expected. *P*-values for amplicon activity versus background were empirically defined using the distribution of standardized residuals (Figure 1B, Supplemental Figure 3E, Supplementary Table 3). We identified 41 amplicons (13%) with increased RNA, suggestive of positive transcriptional regulatory activity. We also compared our linear model directly to RNA/DNA ratio, a common metric used to report activity in MPRAs^13, 32, 33^. We found that 71 amplicons were considered active using the criteria RNA/DNA ratio > 1.5 and RNA/DNA standard deviation < its mean. Amplicons that exhibited RNA/DNA ratio > 1.5 but did not pass the model-based P-value potentially included sequences with weaker enhancer activity. 78% (32/41) amplicons identified using model residuals were also active in the ratiometric comparison. Among 41 positive amplicons with significant regression residual activity (P < 0.05), 20 (49 %) had mean ratio – 1 s.d. > 1.5 across individual replicates, representing the amplicons with the strongest and most consistent MPRA-defined enhancer activity (Figure 2B).

**Figure 2.**
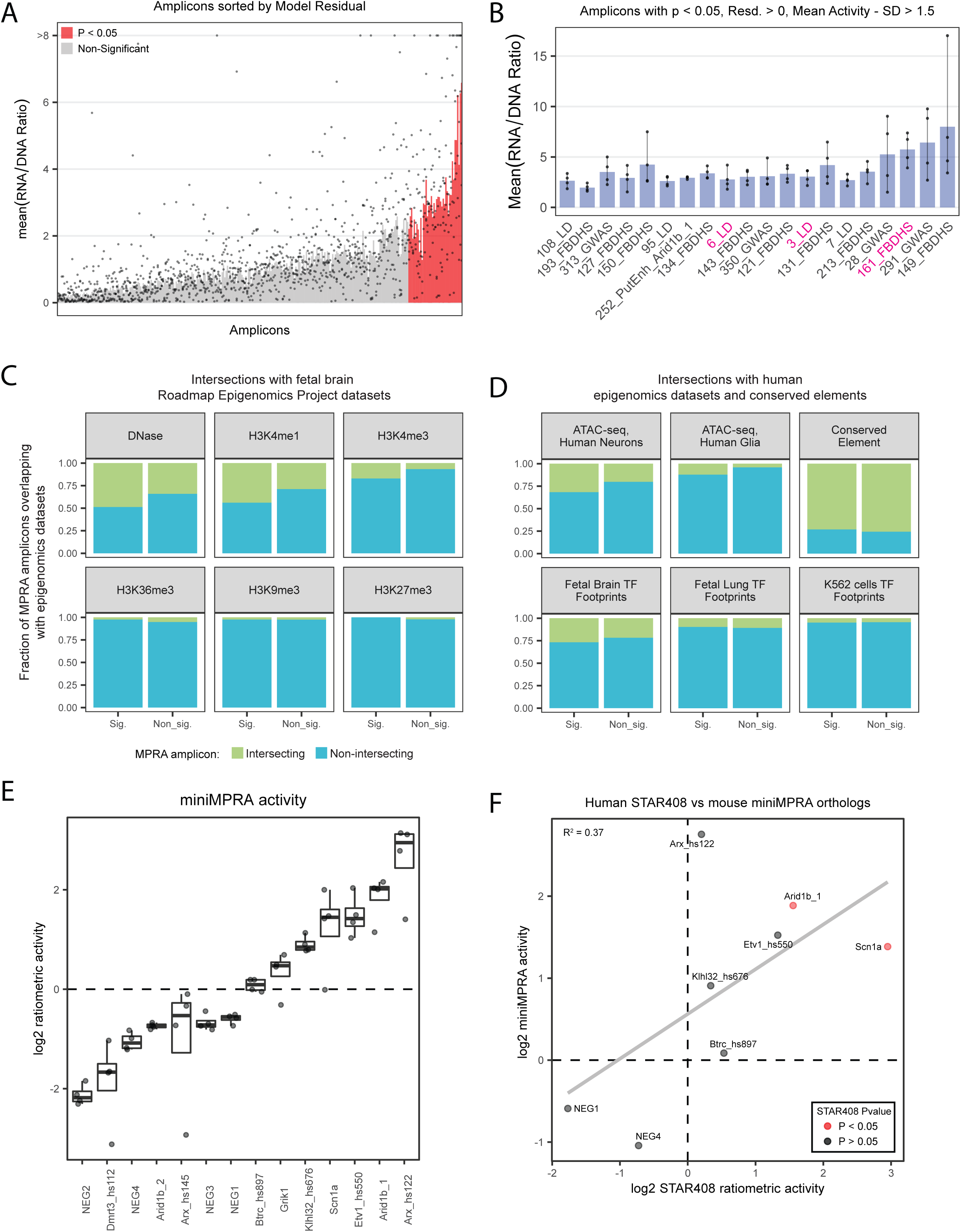
Distribution of active amplicons and concordance with parallel datasets. (**A**) Bar plot representing distribution of activity in the test assay, sorted by linear model residuals. Amplicons are colored as in Figure 1B. To control low-count variability, 1000 counts were added to RNA and DNA counts to stabilize ratios where counts were low. (**B**) The top 20 active amplicons with consistent activity across both linear model and ratiometric model. Three amplicons were used for downstream validation in single-candidate deliveries (magenta). Bar for all plots represents mean RNA/DNA ratio. Each dot represents individual RNA/DNA ratios for each amplicon (4 dots per amplicon). (**C**) Amplicon intersection with fetal brain epigenomic datasets from the Roadmap Epigenomics Project. Amplicons were divided into two groups based on the statistical significance of their activity in the MPRA. (**D**) The same amplicon intersection analysis with various human epigenomics datasets and vertebrate conserved elements, see text for details of intersection datasets. (**E**) Ratiometric activity of miniMPRA mouse library. (**F**) Correlation of the mouse orthologue miniMPRA with the human orthologues of the same sequence from the full MPRA library. Only amplicons that passed quality control criteria for both experiments are included.

We intersected our amplicons with fetal brain epigenomics datasets from the Roadmap Epigenomics Project^34^. We found higher than expected intersection of significant amplicons with genomic loci characterized as DNase hypersensitive (DNase, P = 0.0355). We also observed that significant amplicons had increased enrichment for H3K4me1 (P = 0.0274), and H3K4me3 (P = 0.0131), epigenomic marks associated with transcriptional activity and enhancer function. We did not find enrichment for H3K36me3, a histone mark associated with gene bodies, or for H3K9me3 and H3K27me3, histone marks associated with heterochromatin (Figure 2C).

We also intersected our amplicon activity with a human neuron and glia ATAC-seq dataset from Brain Open Chromatin Atlas (BOCA)^35^, vertebrate evolutionary conserved elements (UCSC Genome Browser), and digital transcription factor (TF) footprints^36^. Similar to Figure 2C, we found marginally significant increased overlap of significant amplicons with human neuron ATAC-seq peaks (P = 0.0497). We also found increased enrichment of significant amplicons in open chromatin in human glia ATAC-seq (P = 0.0159). There were no significant differences in enrichment among amplicon groups in TF footprints in fetal brain or lung, nor in K562 cell lines, nor for conserved elements (Figure 2D). Overall, this enriched overlap between MPRA active candidates and enhancer signatures indicates that activity in our assay has biological relevance.

### Enhancer activity reproduces across MPRA studies

In parallel with this *in vivo* MPRA experiment, we performed a smaller scale parallel reporter (miniMPRA) assay using the same techniques but testing orthologous mouse sequences representing putative enhancer and negative control sequences for our full set (Supplementary Table 4). As human orthologs of these sequences were included in the full MPRA library, this experiment enabled comparison of human and mouse sequences in an *in vivo* mouse brain context and further validation of reproducibility for our MPRA results. We observed robust ratiometric activity in the miniMPRA from candidate enhancer sequences, and we did not observe ratiometric activity in predicted negative sequences (Figure 2E). We also observed a correlation between activity in the mouse miniMPRA and the human orthologs in the full library, indicating that the MPRA activity reflects reproducible enhancer activity and that there is conserved function between orthologous mouse and human sequences tested in the same postnatal mouse brain context (Figure 2F).

### Confirmation of *in vivo* P7 cortex MPRA enhancer results

The STARR-seq orientation places the candidate test sequence in the 3’ UTR of the reporter gene (here EGFP). Enhancers are proposed to function independent of orientation and the position relative to the transcriptional start site^37^, as has been demonstrated in *in vitro* applications of STARR-seq^20, 33^. However, we sought to verify that an enhancer sequence in the STARR-seq orientation not only increased transcription generally, but also did so in a cell-type defined restricted manner *in vivo* when paired with the *Hsp68* minimal promoter and delivered via scAAV. To validate this, we cloned a GABAergic-biased mouse *Dlx5/6* enhancer (*mDlx)*^16, 38^ into the AAV vector used for plasmid library construction. scAAV9 carrying Hsp68-EGFP-*mDlx* was mixed with AAV9 carrying the constitutively active CAG-mRuby3 and this mixture was delivered to P0 mouse forebrain via brain intraventricular injection and collected at P7. The expression of mRuby3 was used as a positive control and to locate the injection site for analysis. Red fluorescent cells, marking the focus of viral exposure, were found distributed in the cortex and hippocampus (Figure 3A, Supplemental Figures 4 and 5A, B). We analyzed EGFP-expressing cells in these regions. The brains were fixed, cryosectioned, and stained with an antibody for GFP to enhance detection of *mDlx* activity (see Methods). We assessed cell-type specificity by counterstaining with an antibody for Lhx6, a cardinal transcription factor expressed in both developing and mature neurons derived from the medial ganglionic eminence^39, 40^.

**Figure 3.**
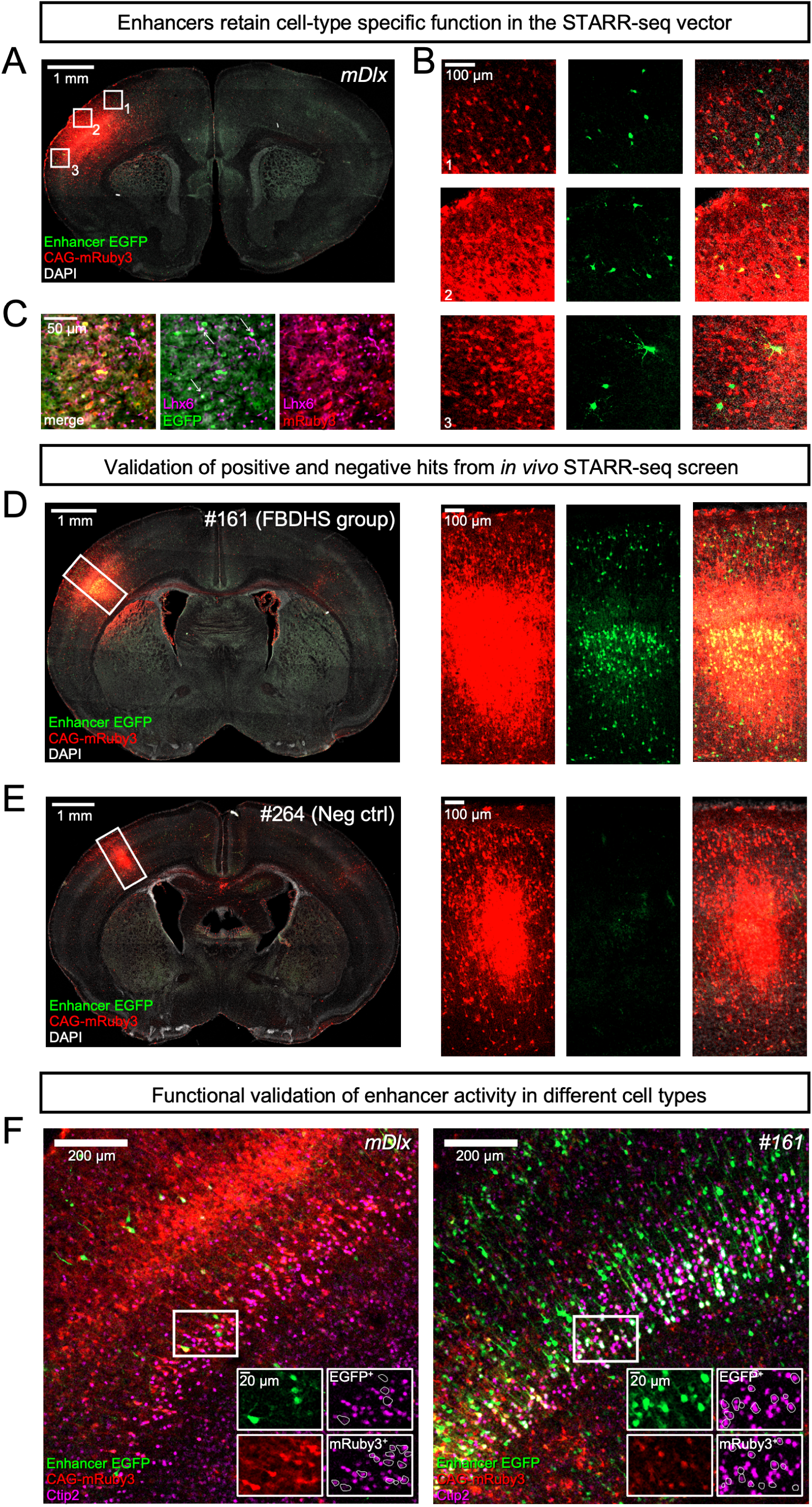
Functional validation of STARR-seq screen. (**A-C**) Enhancers retain cell-type specific function in the STARR-seq vector. (**A**) Representative image of a coronal section of a P7 mouse brain injected at P0 with a virus mixture consisting of an AAV containing the STARR-seq vector carrying the inhibitory interneuron enhancer *mDlx* (scAAV9-Hsp68-EGFP-mDlx) and an injection control AAV containing an expression vector for mRuby3 under the control of CAG, a general mammalian promoter. EGFP expression was visualized via IHC using an anti-GFP antibody, while mRuby3 expression was visualized using native fluorescence. (**B**) Close up of boxed regions in *A* showing morphology of EGFP-expressing cells in the cortex. (**C**) Sections from P7 mouse cortex transduced with *mDlx*-driven STARR-seq reporter vector and mRuby3 injection control at P0, counterstained with an antibody for *Lhx6*, a transcription factor active in deep cortical layer interneurons. EGFP-expressing cells with Lhx6^+^ Nuclei are indicated with arrows. (**D-E**) Validation of positive and negative hits from *in vivo* STARR-seq screen. (**D**) The enhancer candidate amplicon #161, a region overlapping a DNaseI hypersensitive site in fetal human brain and a copy number variant region near *SCN2A*, an autism- and epilepsy-associated gene, which displayed enhancer activity in the STARR-seq screen, was packaged individually into an AAV reporter vector and transduced along with an injection control (CAG-mRuby3) by intracranial injection in P0 mice. A representative image of a coronal section of a transduced P7 brain stained with an anti-GFP antibody is shown (left panel). Close up of the boxed region is shown in the panels on the right (from left to right: Red channel, mRuby3 injection control; Green channel, EGFP expression driven by amplicon #161; Merge with DAPI in grey). (**E**) Representative coronal brain section from P7 mouse transduced as in *D* at P0, but with a reporter vector carrying amplicon #264, a negative control with no predicted enhancer activity that did not display activity in the *in vivo* STARR-seq screen. Close up of boxed region is shown in the panels on the right. (**F**) Functional validation of enhancer activity in different cell types. Enhancers *mDlx* and amplicon #161 were individually packaged into reporter constructs and transduced into neonatal mouse brains with a CAG-driven control virus. Brains were collected at P7 and stained for GFP and Ctip2, a transcription factor necessary for axon development in excitatory projection neurons in Layer V during embryonic development. Inset panels show single channel images (Green, EGFP; Red, mRuby3; Magenta, Ctip2). Ctip2 channel images are shown with EGFP^+^ cells outlined (top) and mRuby3+ cells outlined (bottom). Cells that expressed EGFP under the control of the inhibitory interneuron enhancer *mDlx* displayed a lower frequency of Ctip2^+^ nuclei compared to cells that drove EGFP under the control of amplicon #161 or drove mRuby3 under the control of the general mammalian promoter CAG.

EGFP^+^ cells in these P7 brains had small cell somas and neurites, typical of interneurons compared to pyramidal cells and glia (Figure 3A, B); many were Lhx6^+^ (Figure 3C, Supplemental Figure 4). Since *mDlx*-driven expression of EGFP should be restricted to multiple GABAergic interneurons in the neocortex, we decided to systematically analyze the morphology of EGFP^+^ cells (Supplemental Figure 5A-C). The principal excitatory neurons in the cortex and hippocampus have a typical pyramidal shape with a tear-drop cell body and a long, primary apical dendrite, while inhibitory interneurons are more variable in their appearance. Furthermore, excitatory neurons in the hippocampus are restricted to specific pyramidal and granule layers, and spatial location could be used as a second classification criterion there to lend extra confidence to the morphological assessment. Based on these morphological and spatial criteria and the observation that many express Lhx6, the vast majority of EGFP-expressing cells resemble interneurons (92.7%, n = 314 cells in 5 animals, Supplemental Figure 5A, C). 3.2% of EGFP^+^ cells exhibited pyramidal morphology. We compared these results with expression driven from the same *mDlx* sequence in a conventional orientation upstream of a minimal β-Globin promotor^16^. The specificity for interneurons in our hands was similar in both constructs (Supplemental Figure 5B, C). These data, taken together, indicate that enhancer orientation and choice of minimal promoter did not greatly affect cell type-specific activity of the enhancer reporter construct.

Using the same Hsp68-EGFP 3’-UTR oriented backbone, we next cloned individual enhancers for selected positive and negative hits from the MPRA screen and generated scAAV9 for each construct to validate enhancer activity in the mouse brain at P7. Amplicon #161 (FBDHS group), an enhancer candidate that overlaps both a DNaseI hypersensitive site in fetal human brain and a copy number variant region near the autism- and epilepsy-associated gene *SCN2A*, and that displayed particularly strong activity in the screen (Supplemental Figure 6A), drove consistent expression of EGFP in the mouse brain at P7 (Figure 3D). On the other hand, a predicted negative sequence that did not display enhancer activity in the screen, amplicon #264, did not drive expression of EGFP (Figure 3E, Supplemental Figure 6B).

To further validate enhancer function in excitatory glutamatergic neurons, we counter-stained for Ctip2, a transcription factor involved in axonal development in excitatory cortical projection neurons in Layer V of the cortex^41, 42^. Although expression of Ctip2 is not exclusive to excitatory neurons in adult mice^43^, it is commonly used as an excitatory Layer V marker during embryonic and early postnatal development^42, 44–46^. We reasoned that if our *in vivo* MPRA reporter construct accurately reproduced cell-type specific enhancer activity, then we should see a difference in the Ctip2 overlap with EGFP driven by amplicon #161 compared to EGFP driven by *mDl*x. Indeed, after counterstaining brain sections transduced as described above with antibodies for Ctip2, we found that EGFP^+^ cells driven by *mDlx* exhibited significantly lower frequencies of Ctip2^+^ nuclei compared to either CAG-driven mRuby3^+^ cells or EGFP^+^ cells driven by amplicon #161 (Figure 3F, Supplemental Figure 5D). These results demonstrate that our MPRA could accurately reflect enhancer activity of particular candidate sequences *in vivo*.

### Dissection of regulatory elements within the third intron of *CACNA1C*

Our library included amplicons spanning across a psychiatric disorder-associated LD interval within the ∼330 kb third intron of the gene *CACNA1C*, which encodes the α1 subunit of the L-type voltage-gated calcium channel Ca_V_1.2. This region contains previously *in vitro* defined regulatory elements^47, 48^ harboring schizophrenia- or bipolar disorder-associated SNPs, predominantly *rs1006737*^49, 50^, *rs2007044*^23^, *rs4765905*^51^, and *rs4765913*^23^. Via MPRA, we assessed the activity of 17 amplicons within the *CACNA1C* intron covering SNPs in LD (r^2^ > 0.8). As a comparison set, we also included 5 amplicons covering SNPs in LD (r^2^ > 0.8) associated with a non-neuronal SNP associated with hematocrit (*rs7312105*)^52^ (Supplemental Figure 7).

Three amplicons within the *CACNA1C* psychiatric disorder LD interval drove significant RNA transcript expression in both linear and ratiometric models in our MPRA: amplicons #3 (overlapping *rs1108075* and *rs11062166*), #6 (overlapping *rs12315711* and *rs2159100*), and #7 (overlapping *rs11062170* and *rs4765905*) (Figure 4A). In comparison, no amplicons from the hematocrit LD interval passed significance threshold for activity (Supplemental Figure 7). We validated #3 and #6 (highlighted in blue, Figure 4A) for enhancer activity in postnatal brain using our single-candidate reporter construct strategy. We also validated lack of activity for #2, an amplicon in the same LD block that did not have significant enhancer activity based on the MPRA results (highlighted in red, Figure 4A). Similar to the negative controls and consistent with MPRA findings, #2 did not drive significant EGFP expression in the brains of P7 mice (Figure 4B, top panels). On the other hand, #3 and #6 drove detectable EGFP expression in cells throughout the cerebral cortex wherever viral exposure was detected by the co-injected CAG-driven mRuby3 positive control (Figure 4B, middle and bottom panels respectively). In a follow-up experiment, we transduced scAAV9-Hsp68-EGFP-#3 and AAV9-CAG-mRuby3 at P0 but waited to collect the brains until P28, at which time we observed that amplicon #3 drove EGFP expression in cortical neurons of adolescent mice (Figure 4C, Supplemental Figure 8). These results demonstrate that the AAV MPRA implementation was effective at screening putative regulatory sequences for *in vivo* enhancer activity in the brain, with activity concordant between MPRA results and individual positive and negative EGFP expression as defined via individual delivery and imaging of P7 mouse forebrain, and that these results could be extended to study enhancer activity patterns in later development.

**Figure 4.**
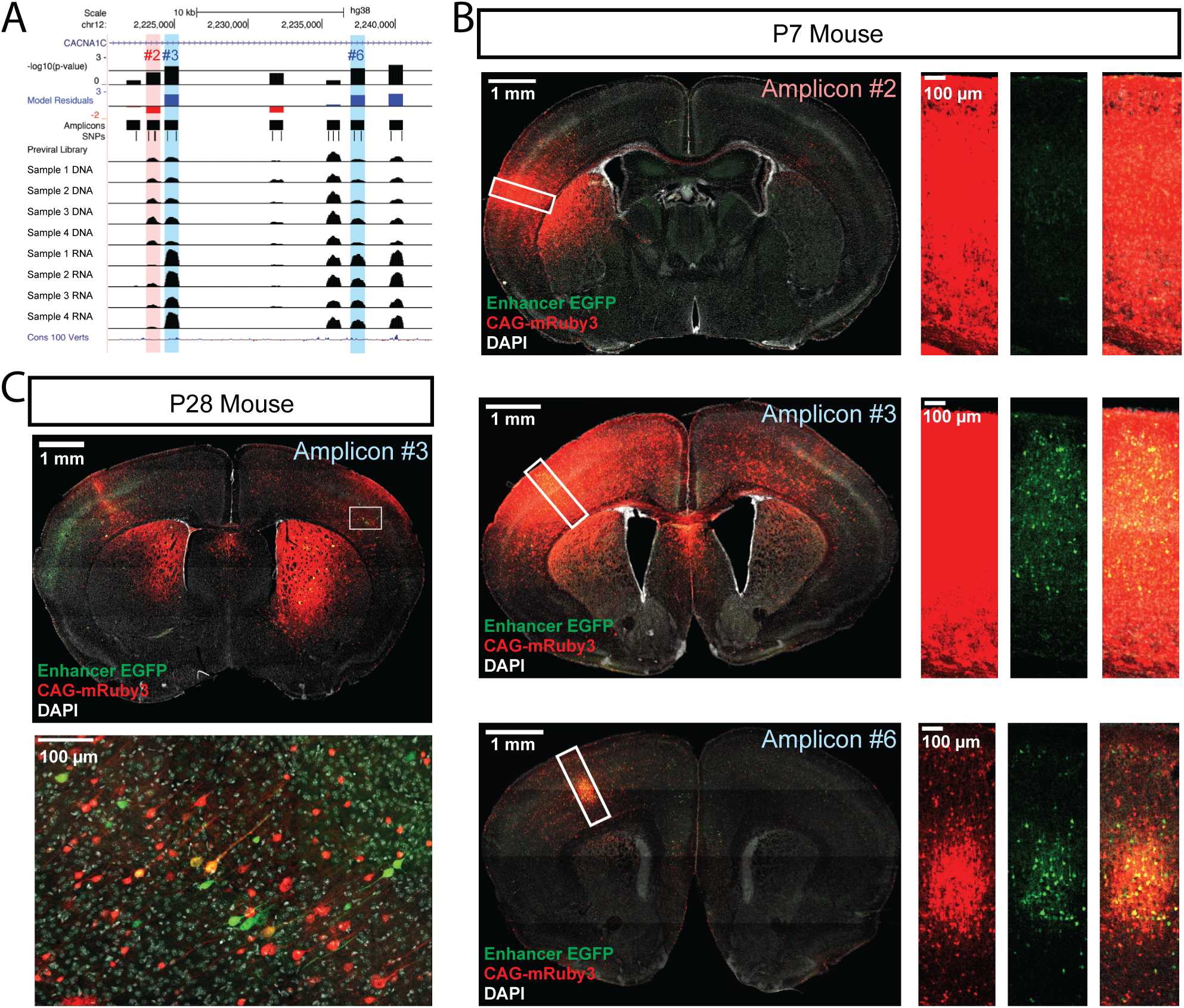
Functional dissection of the large third intron of *CACNA1C*. (**A**) UCSC Genome Browser representation of amplicons #1 through #7 in the third intron of CACNA1C (hg38, chr12:2,220,500-2,242,499). Normalized coverage of aligned reads for DNA and RNA samples are shown for the four biological replicates. Normalized coverage of aligned reads for DNA is also shown for the pre-viral plasmid library. Three amplicons, #3, #6, and #7, were found significantly active in our assay. (**B**) Confocal images of single-candidate validation of amplicons #2, #3, and #6 (red and blue highlighted amplicons in A). Mice were transduced at P0 with two AAV vectors: one for an HSP68-EGFP-3’UTR enhancer reporter construct carrying the indicated amplicon and a second control vector, CAG-mRuby3. Brains were fixed at P7 and sectioned and stained with an antibody for EGFP for signal amplification. Tiled, whole section images are shown on the left. Close up of boxed regions are shown in the panels on the right. Green, EGFP; red mRuby3; grey, DAPI. These experiments validated robust EGFP expression driven by the two positive MPRA hits (#3 and #6), with substantial EGFP reduction for the MPRA negative amplicon #2. (**C**) Mice were transduced with AAV including positive amplicon #3 and processed as in B, but were raised to P28 before fixing, sectioning, and staining.

## DISCUSSION

The ability to test in parallel the regulatory capacity of candidate sequences *in vivo* offers considerable potential for elucidating the role of enhancers in the developing brain and for efficient identification of enhancers that are capable of driving precise expression patterns. Here, we report successful rAAV-mediated delivery of a 3’-oriented parallelized enhancer reporter assay to early postnatal mouse brain and demonstrate the utility of this approach in screening human DNA sequences for regulatory activity in the brain, identifying and validating enhancers in disease-associated loci. Via this MPRA, we identified novel enhancers active in P7 mouse cortex, showcasing example applications including identifying regulatory sequences associated with ASD-associated loci, screening amplicons that include lead SNPs from human genetic studies, and comprehensive testing for enhancers harboring SNPs from disease associated non-coding LD intervals. We show that amplicons active in our MPRA were more likely to have enhancer signature across functional genomics datasets and that orthologous mouse sequences tested in an independent MPRA showed strong activity correlation. Finally, we validated MPRA activity predictions via imaging of EGFP for four positive enhancers and two negative sequences *in vivo* in P7 brain, and we confirmed that activity for one of these positives continued to P28. This study provides a model for applying this powerful screening approach *in vivo* in mammalian brain.

Our study represents one of the first parallelized enhancer assays testing human sequences in early postnatal mouse brain via rAAV-mediated delivery. We validated MPRA performance, verifying both the capacity for the construct to drive characteristic cell-type restricted expression for a known interneuron enhancer, as well demonstrating EGFP protein expression driven by novel putative enhancers that were active *in vivo* in P7 mouse cortex. Based on MPRA and validation experiments, the main sources of variation across MPRA replicates were AAV transduction rate and injection site (Supplemental Figures 9, 10, and 11), highlighting the need for delivery controls to ensure reproducibility when applying MPRAs *in vivo* via viral delivery. We observed some variability in amplicon RNA dropout across replicates, likely due to a combination of transduction efficiency variability across animals, sensitivity recapturing amplicons with low viral representation, and PCR stochasticity^53^. While not an obvious issue here, effect of PCR-based clonal amplification bias in other MPRAs has been shown to be reduced with the addition of barcodes and unique molecular identifiers to the reporter construct^54^.

Amplicons in this assay were approximately ∼900 bp long, representing some of the longer sequences tested in MPRAs to date. Our rationale, that longer sequences afford more native biological context to enhancer activity, appears in accordance with a recent study comparing MPRA designs^33^, which finds that longer sequences add biological signal including enrichment of an RNA polymerase III catalytic subunit, histone-modifying enzymes, and increased transcription factor binding. However, increased length, especially when inserted into the 3’UTR, may contribute to increased variability across replicates^33^, and may drive mRNA degradation^55^ that would inhibit RNA transcript detection in the assay. We detected a number of amplicons that had reduced RNA compared to expected background levels, and these amplicons may be subject to such RNA degradation. Alternatively, these sequences may have silencer or repressive effect on expression, and such sequences have been reported in recent MPRA screens^56^. While artifacts from inserting large sequences into the 3’UTR may have impacted our results, we nonetheless show that orthologous mouse sequences exhibited correlated presence and absence of activity and similarly that amplicons with absence or presence of MPRA activity across replicates consistently reproduced in independent single deliveries to the brain. Further, active amplicons were enriched for DNAse Hypersensitivity Sites, H3K4m1, and H3K4me3 peaks generated from fetal human brain tissue, and with ATAC-seq peaks from FACS-purified neurons and glia from postmortem human brain. Overall, the combination of reproducible MPRA activity, enrichment for neuron-specific enhancer signatures, and activity validation in AAV experiments provide strong evidence that positive results of the MPRA indeed act as enhancers.

Parallelized enhancer assays such as the one reported here have the possibility to become fundamental for assessing active enhancers in the brain and offer great potential for functional dissection of sequence-based enhancer activity. The vast majority of sequence variants associated with genetic risk for neurodevelopmental and neuropsychiatric disorders are found in non-coding regions^57^, many of which are presumed to be located in enhancers^23^. As an example, our assay enabled functional annotation of 17 intronic *CACNA1C* amplicons spanning regions harboring schizophrenia- and bipolar disorder-associated SNPs within strong linkage disequilibrium and identified three regions with enhancer activity in the brain. Notably, we detected no activity in our assay from amplicon #5 (LD group), which spans a region harboring the SNPs rs1006737 and rs2007044, two of the most statistically significant risk SNPs^21^ from GWAS of schizophrenia and bipolar disorder (Figure 5A). Two of the three amplicons we identified as enhancers in P7 brain had prior evidence of enhancer activity in *in vitro* models^47, 48^, indicating our assay can reproducibly detect enhancer activity identified in orthogonal studies. The region spanning amplicon #6 was previously annotated as a putative enhancer in physical proximity to the *CACNA1C* promoter in human induced pluripotent stem cell (hiPSC)-derived neurons^48^. The region spanning amplicon #7 exhibited enhancer activity via luciferase assay in SK-N-SH cells^47^. In addition, #7 was predicted as putative enhancer based on open chromatin signatures in both fetal and postmortem human brain^5, 34, 35^. Finally, we show that activity of one of these *CACNA1C* intronic enhancers continues at P28, highlighting the potential to apply the MPRA and single enhancer AAV testing at later ages. Although our assay was not designed to directly compare the activity of sequence variants, adapting this assay to improve sensitivity and power to evaluate the effects of sequence variation on enhancer activity will be critical in understanding how mutations within the *CACNA1C* intron contributes to altered gene expression.

In summary, our *in vivo* parallelized enhancer reporter assay in P7 mouse brain enabled us to identify novel enhancers active in early postnatal mouse brain and pinpoint potential regulatory regions where disease-associated sequence variation (e.g. GWAS SNPs) may contribute to transcriptional pathology. Our results highlight the opportunities gained by in-depth functional dissection of enhancers in the developing brain via *in vivo* adaption of MPRAs, toward deeper understanding of enhancer biology across neurodevelopment and in the etiology of neuropsychiatric disorders.

## METHODS

### Selection of library candidates

408 candidates were selected for screening based on the following criteria: 22 regions containing *CACNA1C*-associated SNPs (*rs1006737*, *rs2007044*, *rs4765913*, *rs4765905, and rs7312105*) and variants in linkage disequilibrium (r^2^ > 0.8) of listed SNPs; 70 regions containing *SCN1A*-associated SNPs (*rs7587026*, *rs12987787*, *rs11890028*, *rs6732655*, *rs11692675*) and variants in linkage disequilibrium (r^2^ > 0.8); 29 epilepsy-associated SNPs (GCST001662: *rs13026414*, *rs72823592, rs10496964, rs12059546, rs39861, rs2717068, rs771390, rs10030601, rs12720541;* GCST002547: *rs28498976, rs2947349, rs1939012, rs55670112, rs535066, rs111577701*; GCST002141: *rs492146, rs72700966, rs61670327, rs143536437, rs11861787, rs72698613, rs12744221*; GCST000691: *rs346291, rs2601828, rs2172802, rs2841498, rs1490157, rs2475335;* GCST001329: *rs2292096*); 142 regions overlapping schizophrenia-associated SNPs^23^; 129 regions overlapping DNaseI hypersensitive sites identified from fetal human brain^5, 34^ found within autism-associated copy number variants^26, 27^; and 16 human homologs of mouse sequences scored for putative regulatory activity by mouse H3K27ac data^1^. Schizophrenia- and epilepsy-associated SNPs were identified using the NHGRI-EBI GWAS Catalog^24^. SNPs in linkage disequilibrium for query SNPs listed as *CACNA1C*- or *SCN1A*-associated above were identified using HaploReg v3^58^ using LD reported from the 1000 Genomes Project^25^. Full selection metadata for each region of interest be found in Supplementary Table 1.

### Vector preparation and library cloning

We designed 408 PCR amplicons covering the regions of interest described above by customizing a batch primer design pipeline using Primer3^59^ (Supplementary Table 5). All PCR primers were designed using the human genome assembly GRCh37 and the Primer3 human mispriming library *cat_humrep_and_simple.fa*. In brief, amplicons were designed to fit the following criteria with all other parameters set to the Primer3 default: amplicon product size between 880-940 bases; primer size between 18-25 bases with an optimum of 20 bases; primer anneal temperature between 57-61 °C, with an optimum of 60 °C; maximum primer pair annealing temperature difference of 2 °C; and primer GC content between 30-70%. We excluded any primer pairs that contained a SNP within 50 bases of either end of the PCR product. PCR primers were then appended with homology sequences (forward primer: tgtacaagtaacatagtcatttctagaTTAATTAA; reverse primer: gctctagtcgacggtatcgataagcttGGCGCGCC) for Gibson Assembly cloning, up to a total primer length of 60 bases.

To generate the amplicon library, we performed individual PCRs using Phusion Hot Start II High Fidelity DNA Polymerase (Invitrogen #F549S) in 10 µL reaction volumes in 96-well plates with 50 ng of pooled human DNA (1:1 mix of Coriell DNA pool #NA13405 and Coriell DNA pool #NA16600) as the DNA template. 30 cycles of PCR were performed with annealing at 60 °C and 30 s extension time. We then quantified the concentrations of each PCR reaction using Quant-iT PicoGreen dsDNA Assay Kit (Invitrogen #P7589) to calculate equimolar amounts of each PCR product for downstream pooling.

We next linearized the vector pscAAV-Hsp68-EGFP (Supplemental Figure 1) using PacI (NEB #R0547L) and AscI (NEB #R0558L). AAV DNA vectors are prone to recombination at the ITR sites, so we also performed a separate restriction digest using SmaI (NEB #R0141L) using pscAAV-Hsp68-EGFP to check for vector integrity. The PacI/AscI digested vector was run on a 1.0% agarose gel, excised, and purified using the Promega Gel & PCR Cleanup system (#A9282) and quantified using Nanodrop. After concentration quantification of both vector and PCR products, 5 ng of each PCR product was pooled into one tube for purification using the QIAquick PCR purification kit (Qiagen #28106) and eluted into 30 µL Nuclease Free Water. Pooled DNA concentration and quality was then assessed by Nanodrop and Qubit dsDNA High Sensitivity assay kit (Thermo Fisher #Q33230). We also ran the pooled PCR products on a 1.0% agarose gel to verify the expected product sizes (∼1-1.1 kb).

We performed two separate Gibson assemblies (NEB #E2611S) using 100 ng of PacI/AscI- linearized vector and 3-fold molar excess of pooled PCR amplicons in 20 µL reaction volume. We then incubated the Gibson assembly mixtures for 1 hour at 50 °C. For bacterial transformation, 2.0 µL of Gibson assembly reaction was added to 50.0 µL of NEB Stable competent cells (#C3040I), in triplicate for each reaction. Competent cells were heat shocked at 42 °C for 30 seconds, and recovered in 950 µL of SOC media for 1 hour at 37 °C. 100 µL of each transformation replicate was plated onto 100 µg/mL carbenicillin agar plates and grown overnight at 30 °C. 900 µL of cells were transferred directly into 250-500 mL of 100 µg/mL carbenicillin Luria-Bertani broth and grown overnight on a 30 °C shaker at >250 rpm. Agar plates were compared for colony growth, indicating Gibson assembly success. Individual liquid cultures were purified using the Macherey-Nagel Xtra Maxi EF kit (#740424), eluted into 50 mL Nuclease Free Water, and assessed for concentration and purity using Nanodrop. Randomly sampled individual colonies as well as all maxiprep replicates were assessed by Sanger sequencing to confirm assembly at the cloning site within pscAAV-Hsp68-EGFP, then maxiprep replicates were pooled 1:1 by concentration for the final library used for viral packaging.

### Viral packaging for test library

Adeno-associated virus (AAV9(2YF)^29^ was produced using a triple plasmid transfection method in 293T cells^60^. After ultracentrifugation, the interphase between the 54 and 40% iodixanol fraction, and the lower three-quarters of the 40% iodixanol fraction, were extracted and diluted with an equal volume of phosphate-buffered saline (PBS) plus 0.001% Tween 20. Amicon Ultra-15 centrifugal filter units (Millipore, Bedford, MA) were preincubated with 5% Tween in PBS and washed with PBS plus 0.001% Tween. The diluted iodixanol fractions were buffer-exchanged and concentrated to 250 μL in these filter units. Virus was washed three times with 15 mL of sterile PBS plus 0.001% Tween. Vector was then titered for DNase-resistant vector genomes by real-time PCR relative to a standard.

### Injections of MPRA library virus to neonatal mice

C57BL/6 mice from Jackson Laboratories were used for mouse experiments. All procedures were performed in accordance with the ARVO statement for the Use of Animals in Ophthalmic and Vision Research and were approved by the University of California Animal Care and Use Committee (AUP #R200-0913BC). Surgery was performed under anesthesia, and all efforts were made to minimize suffering. Neonatal mice (on a C57BL6/J background) were injected in the prefrontal cortex in 2 separate locations with 0.5 µL of virus. Only the left side of brain was injected. A beveled 34-gauge needle (Hamilton syringes) was used to push through the skull and to inject virus into the brain. Virus was delivered slowly by hand over the course of 1 min. Mice were anesthetized on ice for 3-4 min prior to injection, and after injection mice were heated on a warm pad before being returned to the cage.

### Tissue collection, library preparation, and sequencing

Bulk forebrains were dissected from postnatal day (P)7 mice injected with the viral library at P0. Dissection included whole forebrain, separated into left and right hemispheres, after removing surface tissue and skull. Hemispheres were stored in RNAlater Stabilization Solution (Thermo Fisher #AM7020) for downstream dual DNA and RNA collection. Hemispheres were then physically homogenized and processed using the AllPrep DNA/RNA Mini Kit (Qiagen #80204) to collect total RNA and genomic DNA from the same tissue. RNA was treated on-column with 1 µL RNase-Free DNase (Qiagen #79254). First strand cDNA was synthesized from purified total RNA using the SuperScript VILO cDNA Synthesis Kit (Invitrogen #11755050) with random primers. We amplified the variable regions of the reporter library from cDNA and genomic (g)DNA using PCR with Phusion Hot Start II High Fidelity DNA polymerase (Invitrogen #F549S) using primers flanking the variable region within pscAAV-Hsp68-EGFP (forward primer, Amplicon PCR Primer F: GATCACTCTCGGCATGGAC; reverse primer, Amplicon PCR Primer R: GATGGCTGGCAACTAGAAGG). PCR consisted of 30 cycles (or 35, for technical replicate Sample 4-35) at 60 °C for annealing and 1 min for extension. PCRs were then loaded onto a 1.0% agarose gel, the ∼1-1.1 kb fragment excised, and gel fragment purified using the Promega Gel & PCR Cleanup system (#A9282). Purified gel fragments were then re-purified using the Qiagen MinElute PCR Purification Kit (#28004) to remove residual chemical contamination, and concentration and quality assessed with Nanodrop and the Qubit dsDNA High Sensitivity assay kit (Thermo Fisher #Q33230). Purified amplicons were prepared for sequencing using the Nextera XT DNA Library Preparation Kit (Illumina #FC-131-1001) using 1 ng of purified amplicon library per sample. Quality and concentration of sequencing libraries was assessed using an Agilent BioAnalyzer instrument. Each library was quantified and pooled for sequencing on the Illumina HiSeq 4000 using a paired end 150-bp sequencing strategy (Supplementary Table 6). The miniMPRA was sequenced on a MiSeq instrument since lower coverage was required due to lower library complexity.

### Sequencing alignment

Raw sequencing reads were trimmed to remove Nextera XT adapters using NGmerge (v0.2_dev)^61^ with maximum input quality score set to 41 (-u 41). Trimmed reads were aligned to the human genome (GRCh38) using BWA-MEM (v0.7.17)^62^ with an option marking shorter split hits as secondary (-M) and otherwise standard parameters. As additional quality control, reads were also aligned to the mouse genome (GRCm38) to check for cross-contamination of mouse DNA from the amplicon PCR. Aligned SAM files were processed using Picard tools (v.2.23.3)^53^: aligned reads were sorted by coordinates and converted to BAM files using *SortSam*, optical duplicates were flagged by *MarkDuplicates*, and BAM file indexing was conducted using *BuildBamIndex*. The number of de-duplicated reads within in silico PCR amplicon coordinates was counted using *samtools view -F 0x400 -c* (v1.7)^63^ and reported as the raw amplicon count.

### Bioinformatics analysis with multiple linear regression

Analyses were performed in R (v4.0.2) and RStudio (v1.3.1093) software. Raw amplicon counts were converted to proportion by normalizing raw count values for each amplicon to all amplicon counts in a given sample. A value of 1 was added to the numerator and denominator to avoid dividing by zero. For downstream analysis, mean amplicon proportions for genomic DNA (“DNA”) and cDNA (“RNA”) were calculated across all samples excluding the technical replicate Sample 4-35. RNA/DNA ratio (“ratiometric activity”) was calculated by dividing porportion_RNA_ by proportion_DNA_ for each amplicon per replicate. After proportion conversion, the dataset was cleaned by removing low count amplicons with less than 200 raw counts in any of the DNA samples, and mean amplicon library proportions less than 2^-15^, resulting in 308 amplicons for downstream analysis.

The dataset used for building the linear model was selected by first removing the 10% of amplicons with the highest and 10% with the lowest mean ratiometric activity across the four biological replicates, resulting in 247 amplicons used for training the model. The model was fit using log_2_-transformed mean RNA and log_2_-transformed mean DNA proportions, with amplicon GC content provided as a model covariate. The normality of residuals distribution was confirmed using the Kolmogorov-Smirnov test and a graphical approach. The model was applied to the complete dataset, and p-values were calculated from the Z-scaled distribution of residuals. We compared the linear activity model to using ratiometric activity alone. Mean ratiometric activity and standard deviation was calculated for each amplicon across all biological replicates, excluding the technical replicate Sample 4-35. We compared the mean ratio, mean ratio – 1. s.d., ranked residuals, and p-value for all amplicons passing initial quality control. We also compared our linear model with a Wilcoxon rank sum with 28/41 amplicons also passing significance in this test.

### Intersections with epigenomics datasets

Amplicons were intersected with epigenomics datasets from Roadmap (macs2 peaks filtered at q < 0.05)^34^, with BOCA merged neuron and glia ATAC-seq peaks from postmortem human brain tissues^35^, with DNAse I digestion based digital TF footprints^36^, and with vertebrate evolutionary conserved elements in the human genome from the UCSC Genome Browser. Consolidated peak intervals (genomic ranges) were merged prior to intersecting using R package GenomicRanges reduce function. Intersections were carried out using GenomicRanges findOverlaps function. Significance of enrichment was calculated using a one-tailed permutation test with 20000 random samplings.

### Single-candidate enhancer reporter cloning

To validate selected positive hits, single-candidate enhancer reporter plasmids were constructed. Candidate sequences, amplicons #2, #3, #6 (LD group), #161 (FBDHS group), or #264 (PutEnh group negative control) were cloned by PCR from a pooled sample of human DNA (1:1 mix of Coriell DNA pool #NA13405 and Coriell DNA pool #NA16600) using the same primer sequences as used to clone the amplicons in the combined test library. Each amplicon was then inserted between the PacI/AscI cloning site downstream of EGFP in pscAAV-Hsp68-EGFP, using In-Fusion (Takara Bio #639649) or Gibson Assembly (NEB #E2611S) according to manufacturer’s specifications. In-Fusion or Gibson reaction products were used to transform Stellar competent cells (Takara Bio #636763) via heat shock at 42 °C and ampicillin-resistant clones were selected at 37 °C using LB-agar plates inoculated with carbenicillin. Successful clones were confirmed by PCR and Sanger sequencing, and the integrity of the viral ITR sequences was verified by restriction digest with XmaI before proceeding with AAV packaging.

### Single-candidate enhancer reporter AAV packaging

Adeno-associated viruses for single-candidate enhancer reporter constructs were packaged in-house using a triple-transfection protocol followed by concentration of viral particles from culture media using a table-top centrifuge protocol^64^. Briefly, for each construct, ∼10^6^ AAV293 cells (Agilent #240073) were plated on a 10 cm tissue culture dish in 10 mL of standard DMEM media (Corning #10-013-CV) supplemented with 10% FBS (ThermoFisher #10082139) and 1% penicillin/streptomycin (ThermoFisher #15070063) and maintained in a humidified incubator at 37 °C and 5% CO_2_. After cells reached ∼60% confluency (1-2 days), the cells were passaged and split onto two 10 cm dishes and grown for an additional 1-2 days before passaging again onto two 15 cm dishes in 20 mL of supplemented culture media each. When the cells in 15 cm dishes were 60-80% confluent the triple-transfection protocol was performed. The media was changed for fresh, supplemented DMEM 1 h before transfection and the culture was returned to the incubator. 17 µg of enhancer reporter plasmid (or CAG-driven mRuby3 control plasmid) was mixed with 14.5 µg AAV helper plasmid (gift from Tian lab), 8.4 µg AAV9 rep/cap plasmid (gift from Tian lab), and 2 mL jetPRIME buffer (Polyplus #114-15), vortexed 10 s and incubated at room temperature for 5 min. 80 µL jetPRIME transfection reagent (Polyplus #114-15) was then added to the plasmid/buffer mixture, vortexed, and incubated at room temperature for 10 min. 1 mL of jetPRIME transfection solution was added dropwise to each 15 cm culture dish which was then returned to the incubator overnight. The next day, the transfection media was exchanged for fresh, supplemented DMEM, and the cultures were incubated for an additional 2 days to allow viral particles to accumulate in the media. Approximately 3 days post-transfection, ∼40 mL of media from both 15 cm culture plates was combined in a 50 mL conical centrifuge tube. Cellular debris was pelleted using a table-top centrifuge at 1500 x g for 1 min, and the supernatant was filtered over a 0.22 µm filter (EMD Millipore #SLGP033RS) into a fresh conical tube. The viral particles were precipitated from the media using AAVanced concentration reagent (System Biosciences #AAV100A-1) according to manufacturer’s instructions. Briefly, 10 mL of pre-chilled AAVanced was added to 40 mL filtered media, mixed by pipetting, and then incubated at 4 °C for 24-72 h. After incubation, viral particles were pelleted by centrifugation at 1500 x g for 30 min at 4 °C. The supernatant was removed, the viral pellet was resuspended in 500 µL cold, serum-free DMEM, transferred to a 1.5 mL Eppendorf tube, and centrifuged again at 1500 x g for 3-7 min. At this step, if there was an AAVanced/viral pellet the supernatant was removed by aspiration and the pellet was resuspended in 40-60 µL cold Dulbecco’s PBS (Fisher Scientific #MT21031CV). If there was no viral pellet after this final centrifugation, we used the viral DMEM solution as our final virus prep. The virus solution was then aliquoted, flash frozen on liquid nitrogen, and stored at -80 °C until use. Once thawed, an aliquot was stored at 4 °C.

Viral titer was estimated by qPCR with an MGB-NFQ FAM-labeled probe that recognized GFP (Life Technologies, sequence: 5’-CTTCAAGGAGGACGGCAA-3’) and 20 µg of Hsp68-EGFP vector plasmid as a standard. Viral DNA and standard plasmid DNA were extracted using the QIAGEN DNeasy Blood and Tissue kit (QIAGEN #69504). qPCR reactions of serial-dilutions of the DNA samples were prepared using TaqMan Universal PCR Master Mix (ThermoFisher #4304437) in a 384-well plate and run on an Applied Biosystems QuantStudio Real-time PCR instrument at the UC Davis School of Veterinary Medicine Real-time PCR Core Facility. Viral titers ranged from 6.8 x 10^10^-2.6 x 10^12^ genome copies/mL, and all viruses were normalized to the lowest titer for each experiment.

### Injections of single-candidate enhancer reporter viruses to neonatal mice

Intracranial virus injections were performed on C57BL/6 mice at P0-1 using protocols approved by the UC Davis Institutional Animal Care and Use Committee (Protocol #20506). First, mouse pups were cryo-anesthetized until cessation of pedal withdrawal reflex. Each pup was then injected in each lateral ventricle with 1 µL of a virus mixture consisting of a 2:1 ratio of enhancer reporter virus to CAG-mRuby3 control virus and containing 0.06% Fast Green dye (Grainger #F0099-1G) (2 µL total, ∼1.4 x 10^8^ particles of enhancer reporter virus). The injection sites for each ventricle were located midway between anatomical landmarks lambda and bregma, approximately midway between the eye and the sagittal suture. After injection, the mouse pup was placed in a warmed recovery chamber for 5-10 min before being returned to its parent’s home cage.

### Immunohistochemistry

After 7 or 28 days of virus incubation, the mice were anesthetized under isoflurane and sacrificed by transcardial perfusion with 4% PFA in PBS. The brain was then dissected and placed in 4% PFA overnight to complete fixation. The fixed brain was dehydrated in a 30% sucrose PBS solution for 3-4 days before freezing in O.C.T. medium (VWR # 25608-930). 30-35 µm coronal sections were cut on a cryostat and collected in PBS. Floating sections were stained in 24-well plates using the following protocol. All steps at 4 °C and room temperature were performed with gentle agitation on an orbital shaker (Cole-Parmer product #60-100 RPM). Antigen retrieval was performed in 1x Citrate buffer (pH 6.0; Sigma-Aldrich #C9999-1000ML) at 60 °C for 1 h. After allowing to cool for 20 min, sections were permeabilized for 20 min at room temperature in PBS containing 0.5% Triton X-100. Sections were then blocked for 1 h at room temperature in a blocking solution containing 0.1% Triton X-100 and 5% milk in PBS. The sections were then transferred to blocking solution containing primary antibodies and incubated for 1-3 days at 4°C. After primary antibody incubation, sections were washed 5 times in 0.1% Triton X-100 PBS for 20 min each time. Sections were then transferred to blocking solution containing secondary antibodies and incubated for 45 min to 1 h at room temperature, followed by 3-5 more 20 min washes in 0.1% Triton X-100. Finally, sections were stained with DAPI (ThermoFisher #D1306) at room temperature for 20-30 min and mounted on slides with ProLong Gold Antifade Mountant (ThermoFisher #P36934). Primary antibodies included chicken anti-GFP (ThermoFisher #A10262, 1:1000) and rat anti-Ctip2 (Abcam #ab18465, 1:500), and secondary antibodies included a Donkey anti-chicken AlexaFluor-488 (Jackson ImmunoResearch #703-545-155, 1:500) and a Donkey anti-rat AlexaFluor-647 (Jackson ImmunoResearch #712-605-153, 1:500 or 1:250).

For immuno-fluorescent co-staining of EGFP and Lhx6, a slightly different staining protocol was used. 30 µm coronal brain sections were first mounted onto microscope slides and then incubated at 37 °C for 30 min. Next, the slides were treated to a series of steps to perform antigen retrieval to detect the Lhx6 antigen. These included 8 min shaking in a base solution of 0.5 g NaOH in 50 mL water, followed by 8 min shaking in 0.3% glycine in 1x PBS, and finally 8 min shaking in 0.3% SDS in 1x PBS. The slides were then incubated in a blocking solution of 5% BSA and 0.3% Triton X100 in 1x PBS for 1 h, followed by primary antibody incubation overnight at 4 °C. The next day, slides were washed in 0.3% Triton in 1x PBS 3 times and then incubated in secondary antibodies for 1 h at room temperature. Slides were finally washed 3 more times and coverslips mounted with Vectashield anti-fade medium (Vector labs, H-1000). Primary antibodies included a mouse anti-Lhx6 (Santa Cruz Biotechnologies #sc-271443, 1:200) and a rabbit anti-GFP (ThermoFisher #A11122, 1:2000), and secondary antibodies included a goat anti-mouse AlexaFluor-647 (ThermoFisher #A32728, 1:300) and a goat anti-rabbit AlexaFluor-488 (ThermoFisher #A32731, 1:300).

### Fluorescence microscopy and image analysis

Confocal images were acquired on a Zeiss LSM800 microscope using a 0.25 NA 5x air objective or a 0.8 NA 25x oil objective. Images were acquired using the same settings for laser intensity, PMT gain, and pixel dwell time. For whole-brain section images showing overall injection and virus expression, images were tiled across the section with a 10% overlap and stitched in postprocessing using the Stitching plugin in Fiji64^65^. Brain sections co-labeled for Lhx6 were imaged using a Leica DM2000 fluorescent compound microscope with a mounted DFC3000 G monochrome camera. Images were acquired using a 10x objective which was sufficient to capture the entire neocortical region from white matter to pia. For all images, cells expressing EGFP or mRuby3 were counted and analyzed using Fiji^66^. Images with co-labeling were then scored for EGFP and mRuby3 cells that were double-positive for the co-label. Example images shown have been processed with a 3×3 median filter in Fiji, background has been subtracted from each channel, and the contrast of each channel has been adjusted to the same levels for comparison across images.

## DATA AVAILABILITY

All supplementary information, including links to raw and processed data, can be found at the Nord Lab Resources page (https://nordlab.faculty.ucdavis.edu/resources/). Software can be found at the Nord Lab Git Repository (https://github.com/NordNeurogenomicsLab/).

## ACKNOWLEDGEMENTS

Sequencing was performed at the UC San Francisco and UC Davis DNA cores. We thank John Flannery for advice and providing space in which we performed preliminary experiments. We also thank the lab of Lin Tian at UC Davis for generously gifting us AAV helper and rep/cap plasmids. This work was supported by NIH/NIGMS R35GM119831. L.S.-F. was supported by the UC Davis Floyd and Mary Schwall Fellowship in Medical Research, the UC Davis Emmy Werner and Stanley Jacobsen Fellowship, and by grant number T32-GM008799 from NIGMS-NIH.

## CONTRIBUTIONS

J.T.L. and L.S.-F. are listed as joint first authors, as each led components of the experiments and analysis. J.T.L., L.S.-F. and A.S.N. designed the experiments. Molecular cloning: L.S.-F., J.T.L., T.L.W., L.C.T.B., I.Z., V.H., and S.M. Viral packaging and library delivery: L.C.T.B. and T.W.S. Bioinformatics: L.S.-F., K.C., Y.W., K.J.L. and A.S.N. Library preparation: L.S.-F. and I.Z. Single candidate viral packaging and delivery: J.T.L. and T.L.W. Immunohistochemistry and imaging: J.T.L., T.W.S., C.P.C., E.C., V.H., S.M., H.P., D.S., and D.V. Image processing and analysis: J.T.L., T.W.S., C.P.C., E.C., V.H., S.M., H.P., D.Q., D.S., and D.V. J.T.L., L.S.-F., K.C., L.C.T.B. and A.S.N. drafted the manuscript. All authors contributed to manuscript revisions.

## COMPETING INTERESTS

The authors declare no competing financial interests.

---

**Supplemental Figure 1.**
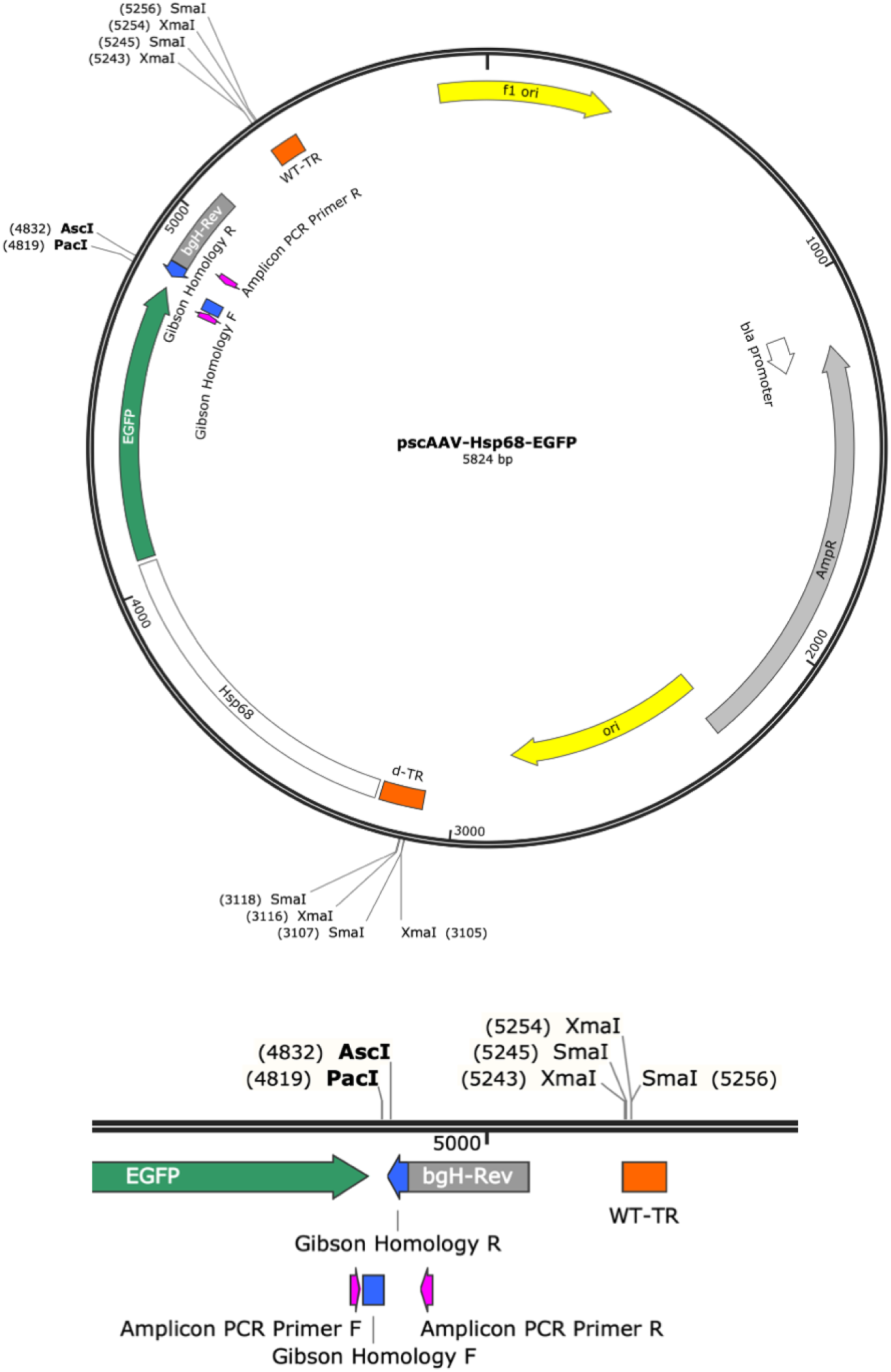
Vector map of pscAAV-Hsp68-EGFP. Vector map of the self-complementary previral vector used for cloning. Inset (bottom) shows multiple cloning site for Gibson assembly and amplicon PCR primer locations.

**Supplemental Figure 2.**
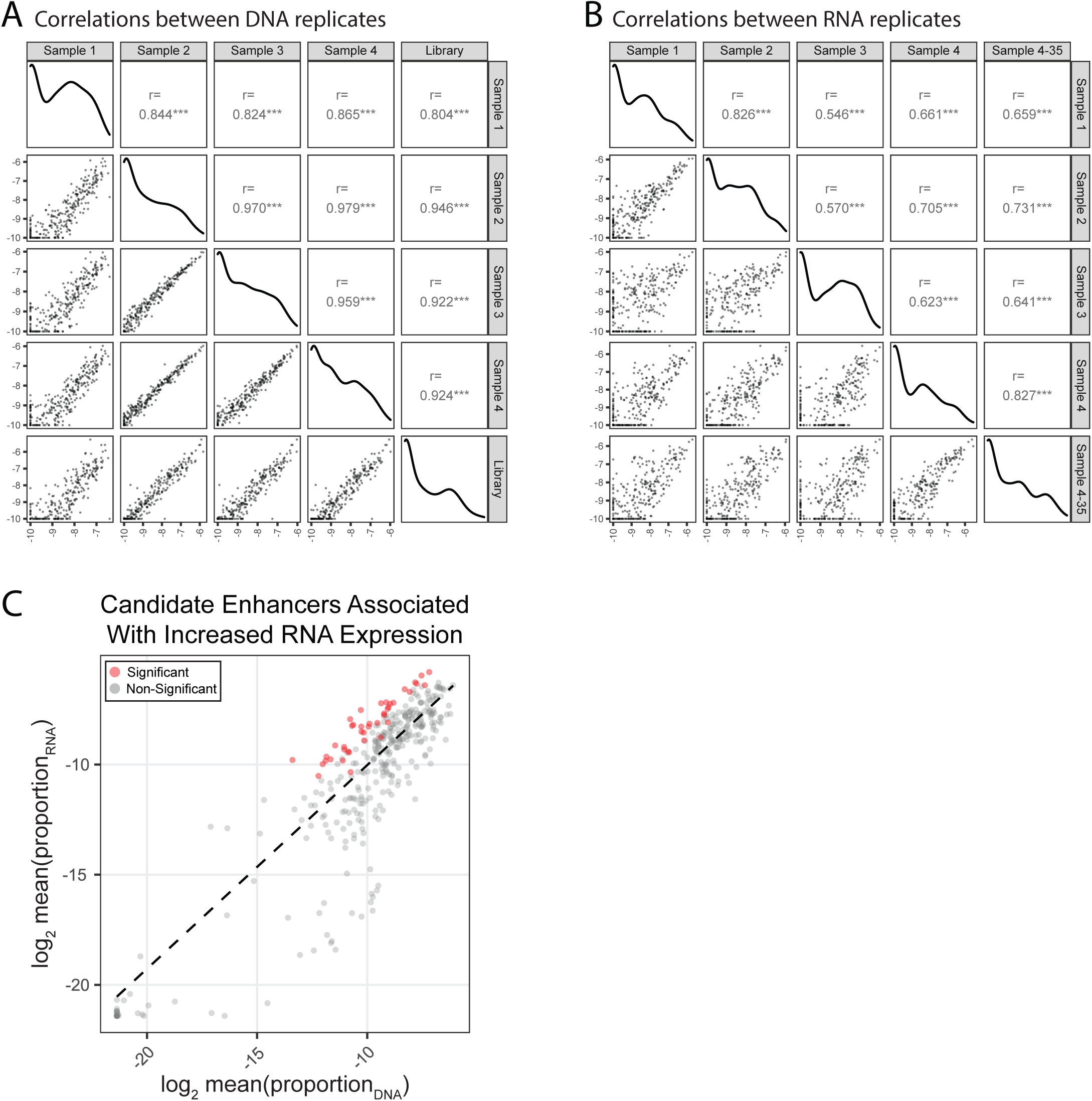
Reproducibility of *in vivo* MPRA. (**A**) Correlation of genomic DNA (“DNA”) representation across biological replicates and the previral plasmid library. Data is shown as log_2_(proportion) per amplicon. Pearson correlation is shown for each pairwise comparison over the filtered dataset (n = 308). (**B**) Correlation of cDNA (“RNA”) representation across biological replicates and a technical replicate of Sample 4 which was subjected to increased PCR cycles during sample preparation (35 cycles instead of the standard 30). Data is shown as log_2_(proportion) per amplicon. Pearson correlation is shown for each pairwise comparison over the filtered dataset (n = 308). (**C**) Correlation of mean RNA/DNA ratio in the assay as shown in Figure 1B, here showing data for all target amplicons (n = 408). Dashed line represents RNA/DNA best-fit line of the data (n = 248). Amplicons are colored by whether they were found significant (P < 0.05) using the multiple linear model.

**Supplemental Figure 3.**
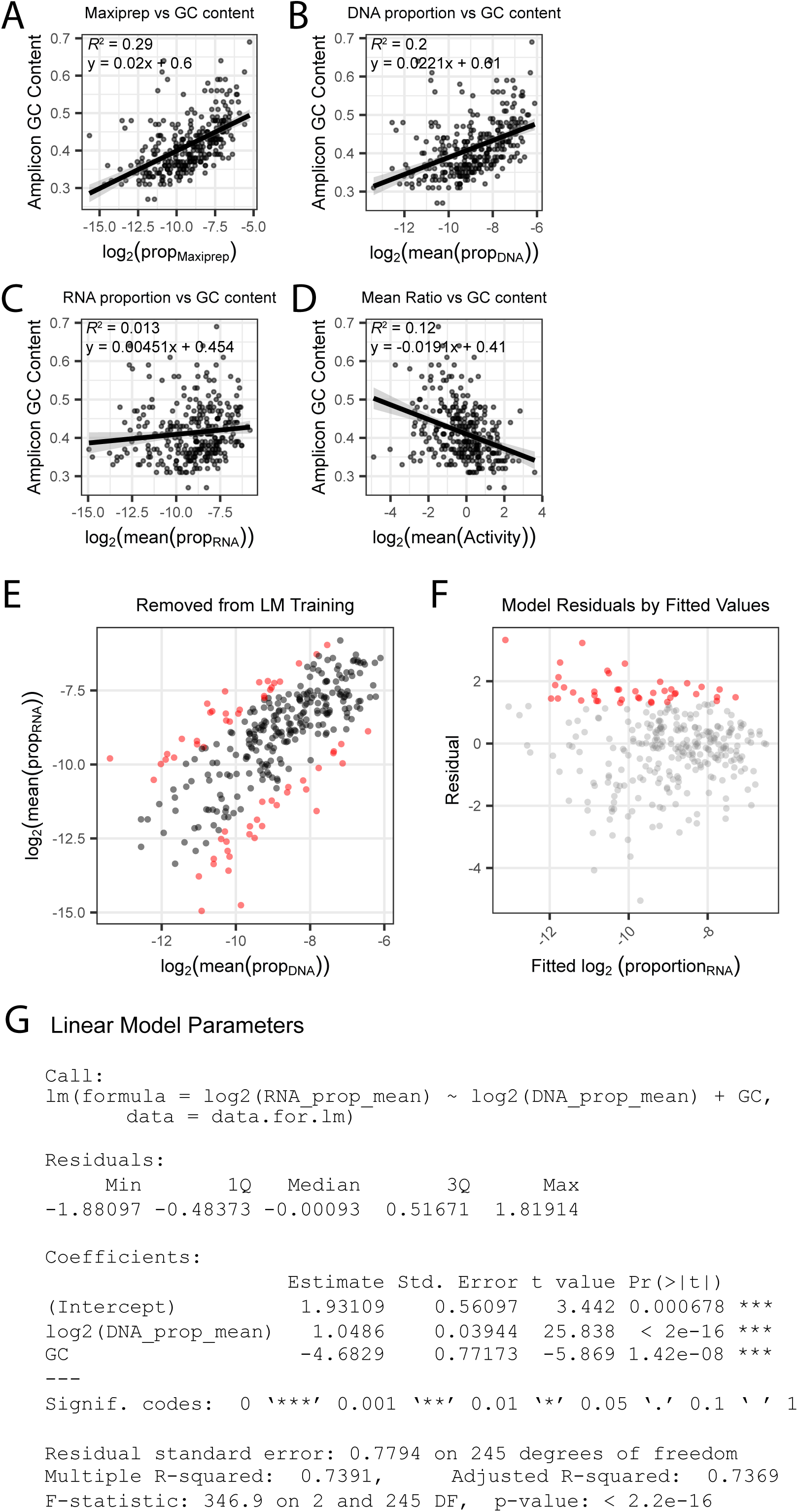
Amplicon selection for linear model building. (**A**) GC content per amplicon by previral library maxiprep proportion. (**B**) GC content per amplicon by mean DNA proportion. (**C**) GC content per amplicon by mean RNA proportion. (**D**) GC content per amplicon by mean RNA/DNA ratio. Line represents linear best-fit line for all plots. Shaded area represents standard error. GC content is correlated with the previral library and the DNA samples, but not to RNA or ratiometric activity. (**E**) Removal of the top 10% and bottom 10% of amplicons, sorted by log_2_ mean RNA/DNA ratio, from linear model building. (**F**) Residuals by fitted log_2_ mean RNA proportion. Amplicons passing significance (p_norm_ < 0.05) are colored red. (**G**) Linear model summary for estimate of background activity.

**Supplemental Figure 4.**
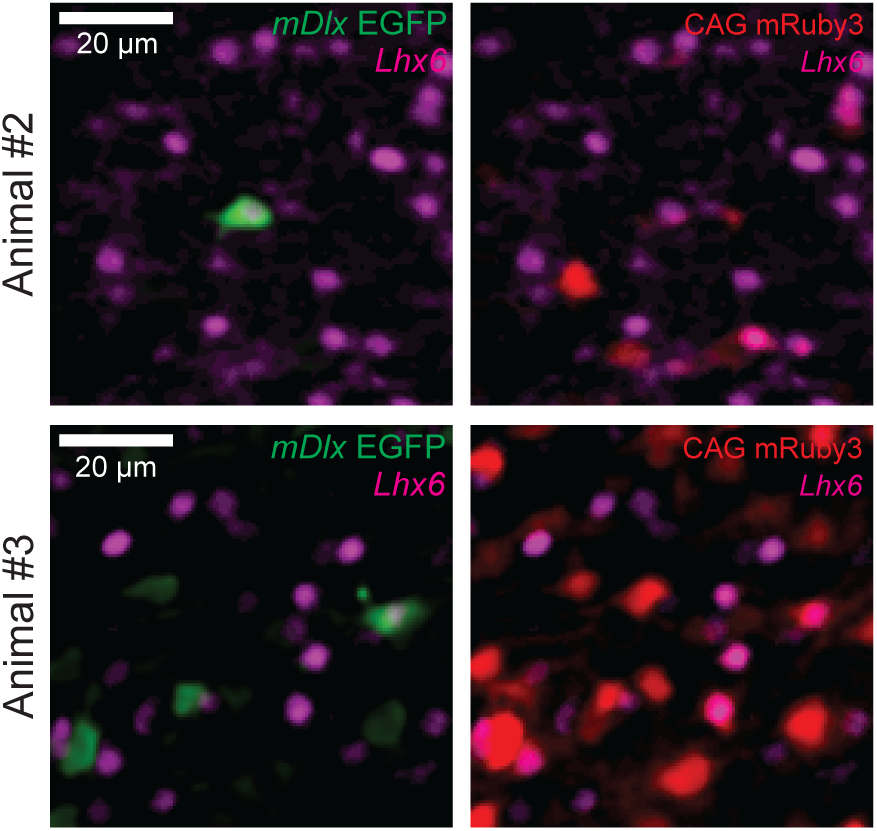
Additional biological replicates of *Lhx6* co-staining of *mDlx-*driven EGFP^+^ cells. Representative fluorescent images of EGFP^+^ cells (left panels) and mRuby3^+^ cells (right panels) in coronal sections of mouse cortex transduced with a mixture of rAAV carrying the constructs Hsp68-EGFP-mDlx and CAG-mRuby3 and counter-stained with an antibody for *Lhx6* (2 additional independent replicates of the representative images shown in Figure 3C).

**Supplemental Figure 5.**
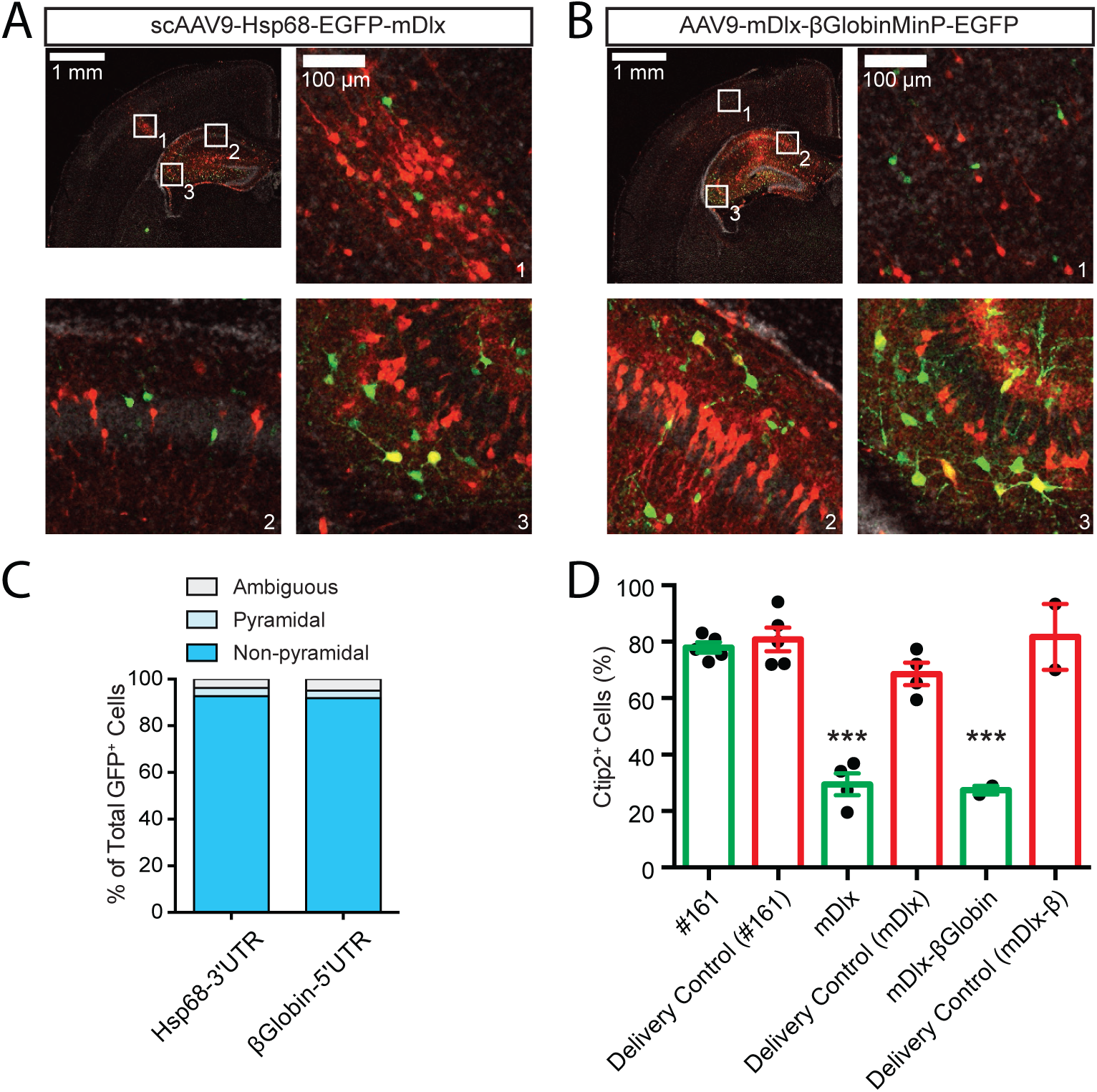
Validation of cell-type specific enhancer function in the STARR-seq orientation. (**A, B**) Representative confocal images of coronal sections of P7 mouse brain transduced by intracranial injection at P0 with scAAV9-Hsp68-EGFP-mDlx (**A**) or AAV9-mDlx-βGlobinMinP-EGFP (**B**) and a CAG-mRuby3 positive control. Sections were stained with an antibody for EGFP for signal amplification. Green, EGFP; red, mRuby3; grey, DAPI. (**C**) Quantification of the numbers of Pyramidal (light blue), Non-pyramidal (darker blue), and cells of ambiguous morphology (grey) observed in confocal imaging experiment in A and B above; N = 5 animals, 314 cells transduced with Hsp68-EGFP-mDlx (Hsp68-3’UTR) and 4 animals, 972 cells transduced with mDlx-βGlobinMinP-EGFP (βGlobin-5’UTR). (**D**) Quantification of images from experiment shown in Figure 4F, with mDlx-βGlobinMinP-EGFP included for comparison. Individual GFP+ and mRuby3+ cells were counted and scored for whether each cell contained a Ctip2-positive nucleus. Cell counts were summed across images for the same brain. Data is presented as mean proportion of Ctip2-positive cells (n = 5 animals co-injected with Hsp68-EGFP-#161 and CAG-mRuby3, 4 animals co-injected with Hsp68-EGFP-mDlx and CAG- mRuby3, and 2 animals co-injected with mDlx-βGlobinMinP-EGFP and CAG-mRuby3).

**Supplemental Figure 6.**
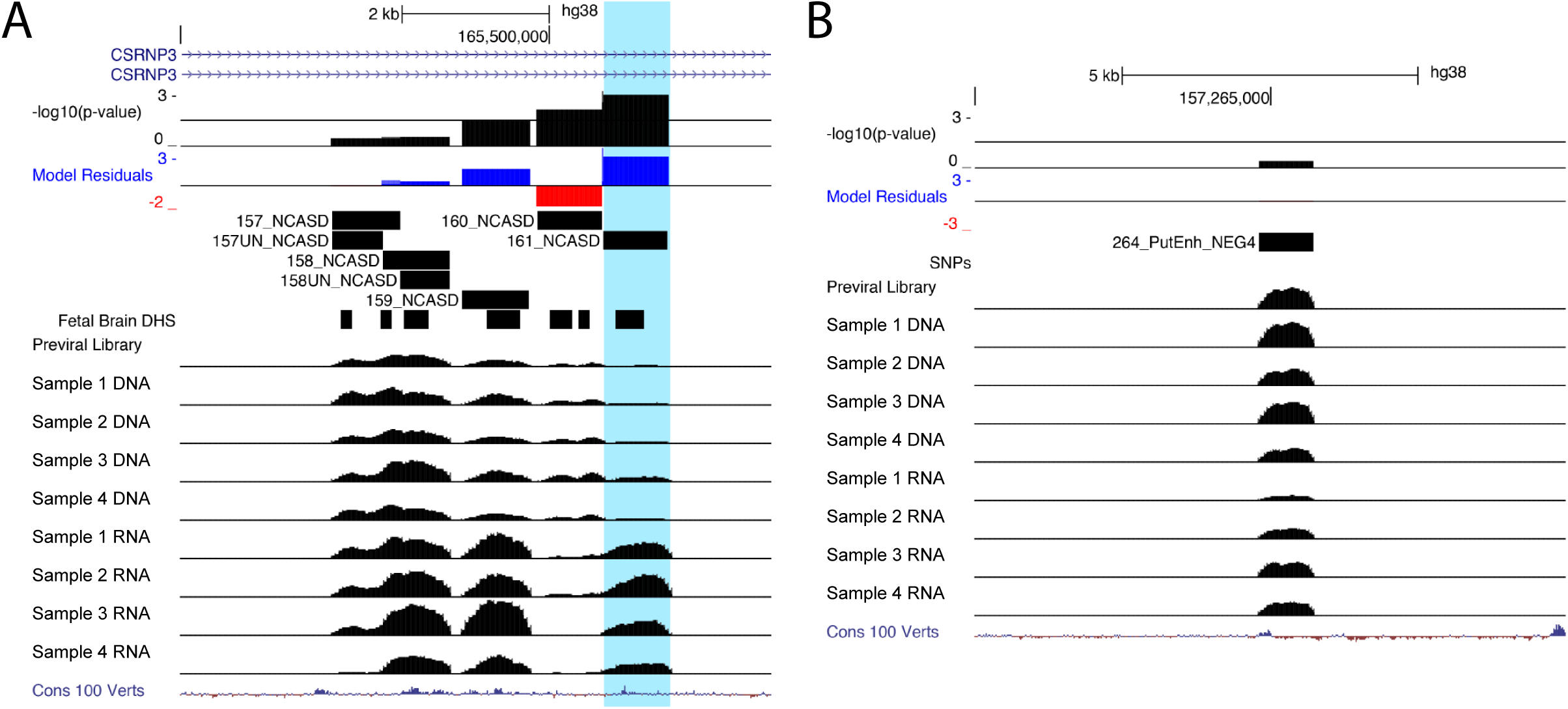
Genome coverage of single candidate validation. UCSC Genome Browser representation of amplicons #161 (**A**), hg38, chr2:165500720-165501601) and #264 (**B**), hg38, chr6:157264812-157265726). Normalized coverage of aligned reads for DNA and RNA samples are shown for the four biological replicates. Amplicon #161 exhibits statistically significant enhancer activity while amplicon #264 does not.

**Supplemental Figure 7.**
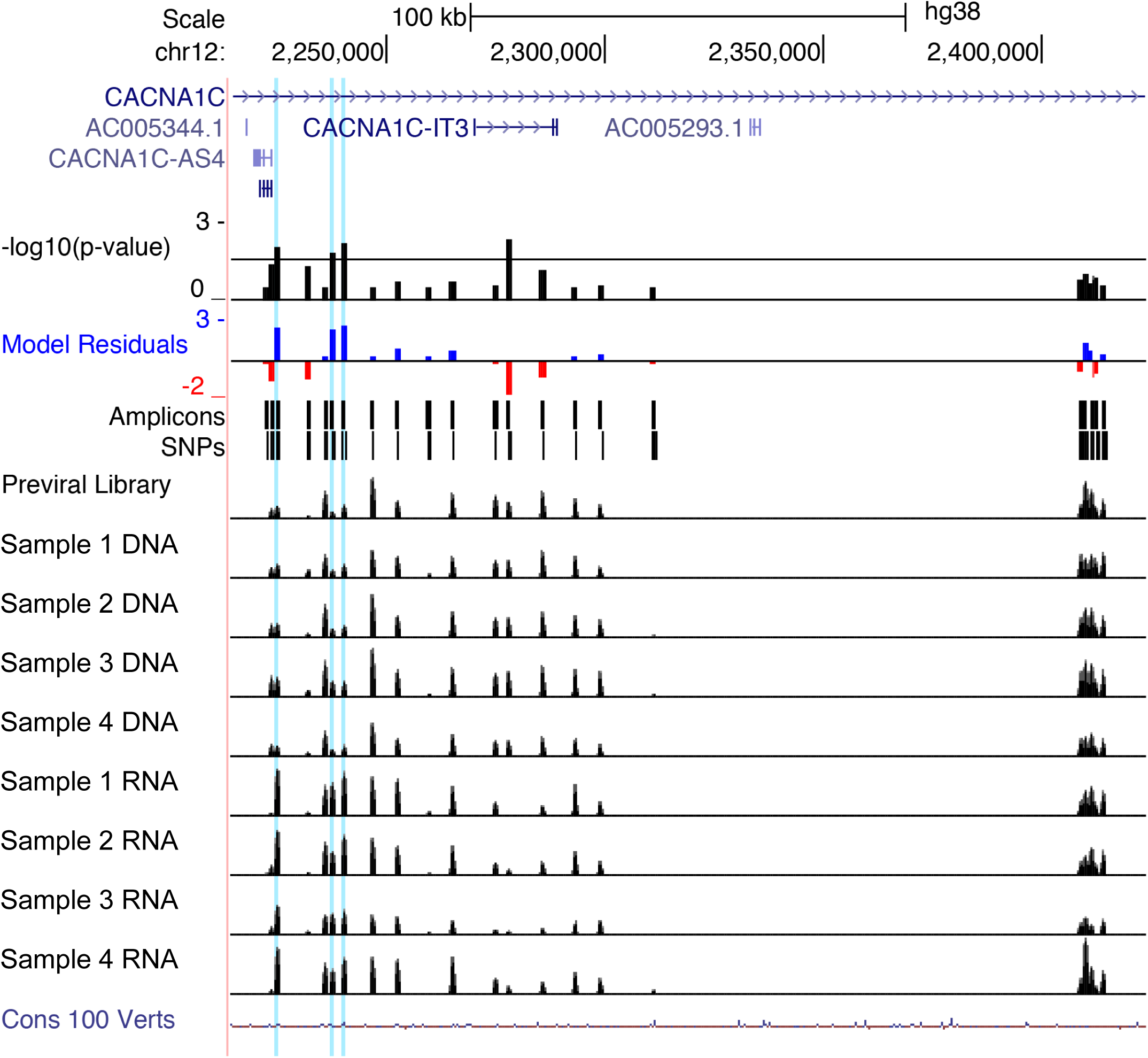
Functional dissection of the third intron of *CACNA1C*. UCSC Genome Browser representation of amplicons #1 through #22 in the third intron of *CACNA1C* (hg38, chr12:2,214,000-2,423,999). Amplicons #18 through #22 (far right) represent regions spanning hematocrit-associated SNPs, which do not show activity in our assay. Normalized coverage of aligned reads for DNA and RNA samples are shown for the four biological replicates. The three amplicons highlighted in blue are #3, #6, and #7, the three *CACNA1C*-associated amplicons with enhancer activity in our assay.

**Supplemental Figure 9.**
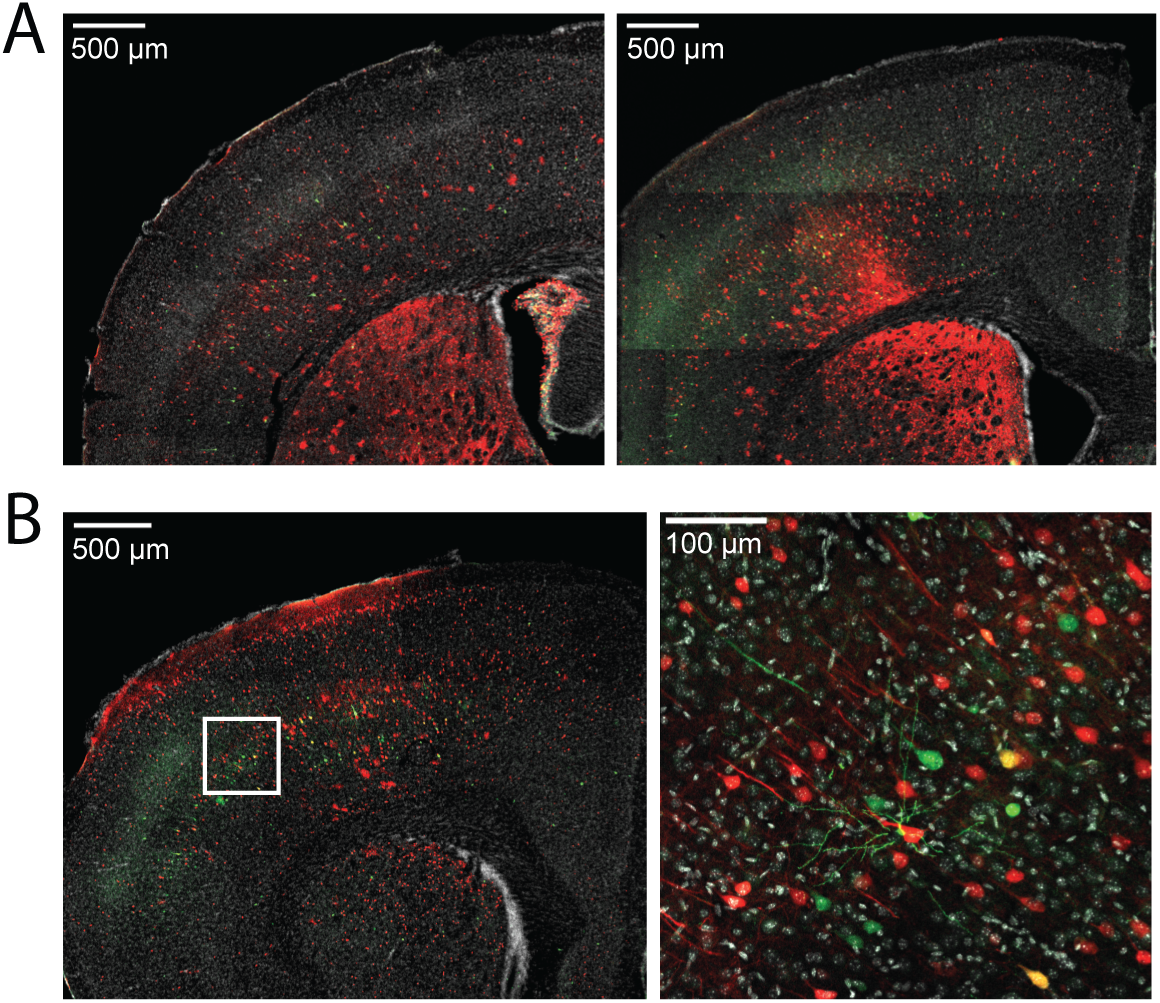
Amplicon #3 in the large intron of *CACNA1C* continues to drive EGFP expression in mouse brain at P28. Mice injected intracranially with scAAV9-Hsp68-EGFP-#3 and AAV9-CAG-mRuby3 at P0 show robust expression of EGFP driven by the enhancer candidate Amplicon #3 at P28. (**A**) 5x confocal images showing EGFP and mRuby3 expression in two representative mice. (**B**) 5x and 25x images showing expression in a third representative mouse.

**Supplemental Figure 9.**
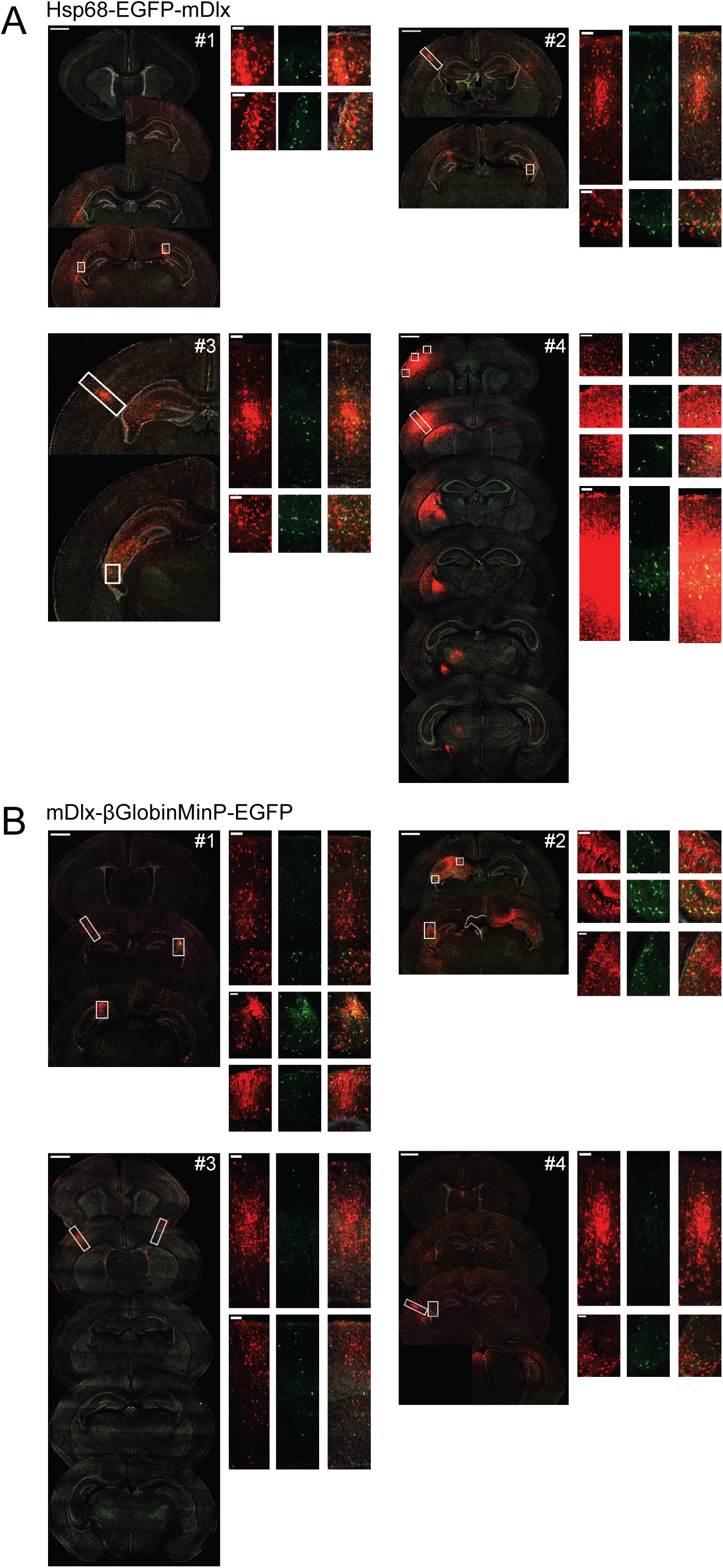
Interneuronal expression of mDlx-driven EGFP expression is consistent despite variable transductions. Representative replicates (4 animals each) of coronal sections from mice transduced at P0 with AAV9-CAG-mRuby3 and either scAAV9-Hsp68-EGFP-mDlx (**A**) or AAV9-mDlx-βGlobinMinP-EGFP (**B**). For each brain, the left-hand panel shows a spread of coronal sections sampling the injection site as it appears across multiple sections of the brain (scale bar = 1 mm), and the right-hand panels are close up views of the boxed regions on the left panel (scale bar = 100 µm). mRuby3 is shown in red, EGFP is shown in green, DAPI is shown in grey.

**Supplemental Figure 10.**
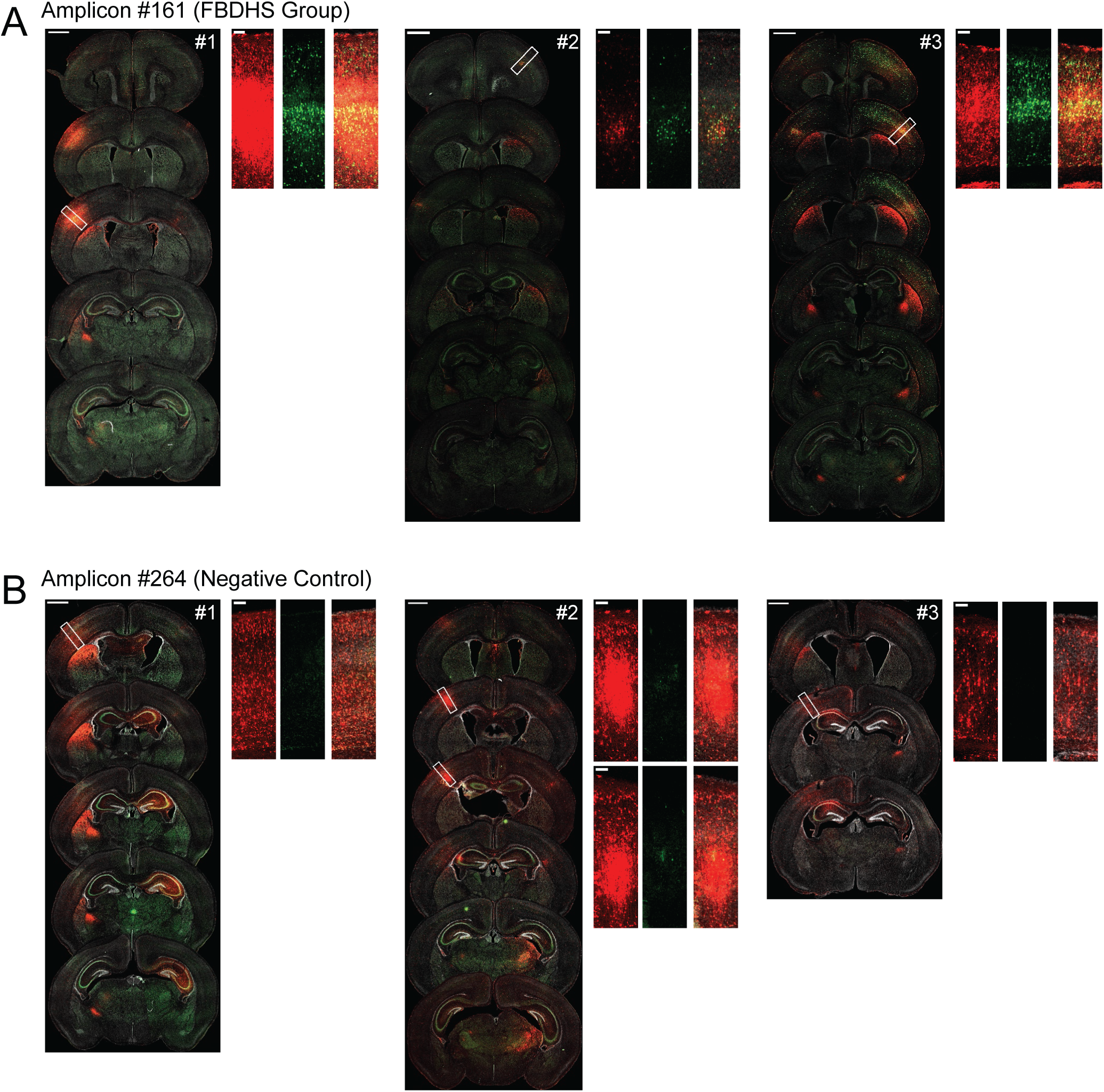
Replicates for validation of positive and negative MPRA hits. Representative confocal images of coronal sections of brains transduced at P0 with AAV9-CAG- mRuby3 and either scAAV9-Hsp68-EGFP-#161 or scAAV9-Hsp68-EGFP-#264, 3 replicates for each condition. For each brain, the left-hand panel shows a spread of coronal sections sampling the injection site as it appears across multiple sections of the brain (scale bar = 1 mm), and the right-hand panels are close up views of the boxed regions on the left panel (scale bar = 100 µm). mRuby3 is shown in red, EGFP is shown in green, DAPI is shown in grey.

**Supplemental Figure 11.**
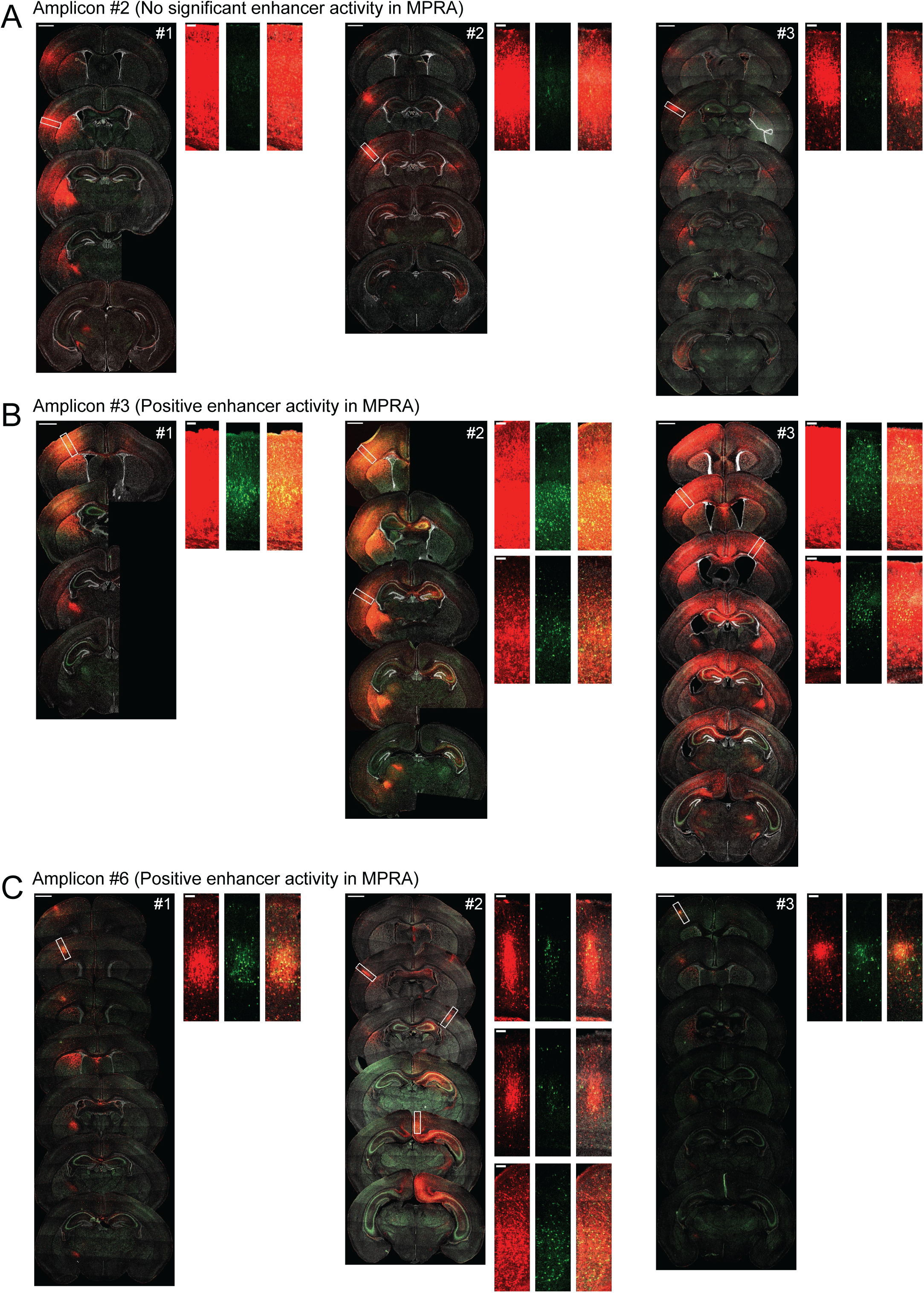
Replicates for functional dissection of *CACNA1C* LD interval. Representative confocal images of coronal sections of brains transduced at P0 with AAV9-CAG- mRuby3 and either scAAV9-Hsp68-EGFP-#2, scAAV9-Hsp68-EGFP-#3, or scAAV9-Hsp68- EGFP-#6, 3 replicates for each condition. For each brain, the left-hand panel shows a spread of coronal sections sampling the injection site as it appears across multiple sections of the brain (scale bar = 1 mm), and the right-hand panels are close up views of the boxed regions on the left panel (scale bar = 100 µm). mRuby3 is shown in red, EGFP is shown in green, DAPI is shown in grey.

## Notes

### Competing Interest Statement

The authors have declared no competing interest.

## REFERENCES

1. Nord, A. S. Rapid and pervasive changes in genome-wide enhancer usage during mammalian development. Cell 155, 1521–1531 (2013).

2. Fuxman Bass, J. I. et al. Human Gene-Centered Transcription Factor Networks for Enhancers and Disease Variants. Cell 161, 661–673 (2015).

3. Calo, E. & Wysocka, J. Modification of Enhancer Chromatin: What, How, and Why? Mol. Cell 49, 825–837 (2013).

4. McLean, C. Y. et al. Human-specific loss of regulatory DNA and the evolution of human-specific traits. Nature 471, 216–219 (2011).

5. Dunham, I. et al. An integrated encyclopedia of DNA elements in the human genome. Nature 489, 57–74 (2012).

6. Moreau, P. et al. The SV40 72 base repair repeat has a striking effect on gene expression both in SV40 and other chimeric recombinants. Nucleic Acids Res. 9, 6047–6068 (1981).

7. Banerji, J., Rusconi, S. & Schaffner, W. Expression of a β-globin gene is enhanced by remote SV40 DNA sequences. Cell 27, 299–308 (1981).

8. Visel, A. et al. A High-Resolution Enhancer Atlas of the Developing Telencephalon. Cell 152, 895–908 (2013).

9. Schoenfelder, S. & Fraser, P. Long-range enhancer–promoter contacts in gene expression control. Nat. Rev. Genet. 20, 437–455 (2019).

10. Benton, M. L., Talipineni, S. C., Kostka, D. & Capra, J. A. Genome-wide enhancer annotations differ significantly in genomic distribution, evolution, and function. BMC Genomics 20, 511 (2019).

11. Kvon, E. Z. Using transgenic reporter assays to functionally characterize enhancers in animals. Genomics 106, 185–192 (2015).

12. Silberberg, S. N. et al. Subpallial Enhancer Transgenic Lines: a Data and Tool Resource to Study Transcriptional Regulation of GABAergic Cell Fate. Neuron 92, 59–74 (2016).

13. Inoue, F. & Ahituv, N. Decoding enhancers using massively parallel reporter assays. Genomics 106, 159–164 (2015).

14. Shen, S. Q. et al. Massively parallel cis-regulatory analysis in the mammalian central nervous system. Genome Res. 26, 238–255 (2016).

15. Shen, S. Q. et al. A candidate causal variant underlying both higher intelligence and increased risk of bipolar disorder. bioRxiv 580258 (2019) doi:10.1101/580258.

16. Dimidschstein, J. et al. A viral strategy for targeting and manipulating interneurons across vertebrate species. Nat. Neurosci. 19, 1743–1749 (2016).

17. Hrvatin, S. et al. A scalable platform for the development of cell-type-specific viral drivers. eLife 8, e48089 (2019).

18. Nord, A. S. & West, A. E. Neurobiological functions of transcriptional enhancers. Nat. Neurosci. 23, 5–14 (2020).

19. Vogt, D. et al. Lhx6 Directly Regulates Arx and CXCR7 to Determine Cortical Interneuron Fate and Laminar Position. Neuron 82, 350–364 (2014).

20. Arnold, C. D. et al. Genome-Wide Quantitative Enhancer Activity Maps Identified by STARR-seq. Science 339, 1074–1077 (2013).

21. Moon, A. L., Haan, N., Wilkinson, L. S., Thomas, K. L. & Hall, J. CACNA1C: Association With Psychiatric Disorders, Behavior, and Neurogenesis. Schizophr. Bull. 44, 958–965 (2018).

22. Abou-Khalil, B. et al. Genome-wide mega-analysis identifies 16 loci and highlights diverse biological mechanisms in the common epilepsies. Nat. Commun. 9, 5269 (2018).

23. Ripke, S. et al. Biological insights from 108 schizophrenia-associated genetic loci. Nature 511, 421–427 (2014).

24. Buniello, A. et al. The NHGRI-EBI GWAS Catalog of published genome-wide association studies, targeted arrays and summary statistics 2019. Nucleic Acids Res. 47, D1005–D1012 (2019).

25. Auton, A. et al. A global reference for human genetic variation. Nature 526, 68–74 (2015).

26. Turner, T. N. et al. Genome Sequencing of Autism-Affected Families Reveals Disruption of Putative Noncoding Regulatory DNA. Am. J. Hum. Genet. 98, 58–74 (2016).

27. Turner, T. N. et al. Genomic Patterns of De Novo Mutation in Simplex Autism. Cell 171, 710–722.e12 (2017).

28. Bevilacqua, A., Kinnunen, L. H., Bevilacqua, S. & Mangia, F. Stage-Specific Regulation of Murine Hsp68 Gene Promoter in Preimplantation Mouse Embryos. Dev. Biol. 170, 467–478 (1995).

29. Dalkara, D. et al. Enhanced gene delivery to the neonatal retina through systemic administration of tyrosine-mutated AAV9. Gene Ther. 19, 176–181 (2012).

30. McCarty, D. M. Self-complementary AAV Vectors; Advances and Applications. Mol. Ther. 16, 1648–1656 (2008).

31. Lee, D. et al. STARRPeaker: Uniform processing and accurate identification of STARR-seq active regions. bioRxiv 694869 (2020) doi:10.1101/694869.

32. Ashuach, T. et al. MPRAnalyze: statistical framework for massively parallel reporter assays. Genome Biol. 20, 183 (2019).

33. Klein, J. C. et al. A systematic evaluation of the design and context dependencies of massively parallel reporter assays. Nat. Methods 1–9 (2020) doi:10.1038/s41592-020-0965-y.

34. Bernstein, B. E. et al. The NIH Roadmap Epigenomics Mapping Consortium. Nat. Biotechnol. 28, 1045–1048 (2010).

35. Fullard, J. F. et al. An atlas of chromatin accessibility in the adult human brain. Genome Res. 28, 1243–1252 (2018).

36. Vierstra, J. et al. Global reference mapping of human transcription factor footprints. Nature 583, 729–736 (2020).

37. Serfling, E., Jasin, M. & Schaffner, W. Enhancers and eukaryotic gene transcription. Trends Genet. 1, 224–230 (1985).

38. Lee, A. T., Vogt, D., Rubenstein, J. L. & Sohal, V. S. A Class of GABAergic Neurons in the Prefrontal Cortex Sends Long-Range Projections to the Nucleus Accumbens and Elicits Acute Avoidance Behavior. J. Neurosci. 34, 11519–11525 (2014).

39. Sandberg, M. et al. Transcriptional Networks Controlled by NKX2-1 in the Development of Forebrain GABAergic Neurons. Neuron 91, 1260–1275 (2016).

40. Alifragis, P., Liapi, A. & Parnavelas, J. G. Lhx6 regulates the migration of cortical interneurons from the ventral telencephalon but does not specify their GABA phenotype. J. Neurosci. Off. J. Soc. Neurosci. 24, 5643–5648 (2004).

41. Arlotta, P., Molyneaux, B. J., Jabaudon, D., Yoshida, Y. & Macklis, J. D. Ctip2 Controls the Differentiation of Medium Spiny Neurons and the Establishment of the Cellular Architecture of the Striatum. J. Neurosci. 28, 622–632 (2008).

42. Leyva-Díaz, E. & López-Bendito, G. In and out from the cortex: Development of major forebrain connections. Neuroscience 254, 26–44 (2013).

43. Nikouei, K., Muñoz-Manchado, A. B. & Hjerling-Leffler, J. BCL11B/CTIP2 is highly expressed in GABAergic interneurons of the mouse somatosensory cortex. J. Chem. Neuroanat. 71, 1–5 (2016).

44. Alcamo, E. A. et al. Satb2 Regulates Callosal Projection Neuron Identity in the Developing Cerebral Cortex. Neuron 57, 364–377 (2008).

45. Leone, D. P. et al. Satb2 Regulates the Differentiation of Both Callosal and Subcerebral Projection Neurons in the Developing Cerebral Cortex. Cereb. Cortex 25, 3406–3419 (2015).

46. Gompers, A. L. et al. Germline Chd8 haploinsufficiency alters brain development in mouse. Nat. Neurosci. 20, 1062–1073 (2017).

47. Eckart, N. et al. Functional Characterization of Schizophrenia-Associated Variation in CACNA1C. PLOS ONE 11, e0157086 (2016).

48. Roussos, P. et al. A Role for Noncoding Variation in Schizophrenia. Cell Rep. 9, 1417–1429 (2014).

49. Ferreira, M. A. R. et al. Collaborative genome-wide association analysis supports a role for ANK3 and CACNA1C in bipolar disorder. Nat. Genet. 40, 1056–1058 (2008).

50. Sklar, P. et al. Whole-genome association study of bipolar disorder. Mol. Psychiatry 13, 558–569 (2008).

51. Hamshere, M. L. et al. Genome-wide significant associations in schizophrenia to ITIH3/4, CACNA1C and SDCCAG8, and extensive replication of associations reported by the Schizophrenia PGC. Mol. Psychiatry 18, 708–712 (2013).

52. van der Harst, P. et al. Seventy-five genetic loci influencing the human red blood cell. Nature 492, 369–375 (2012).

53. Kebschull, J. M. & Zador, A. M. Sources of PCR-induced distortions in high-throughput sequencing data sets. Nucleic Acids Res. 43, e143–e143 (2015).

54. Neumayr, C., Pagani, M., Stark, A. & Arnold, C. D. STARR-seq and UMI-STARR-seq: Assessing Enhancer Activities for Genome-Wide-, High-, and Low-Complexity Candidate Libraries. Curr. Protoc. Mol. Biol. 128, e105 (2019).

55. Rabani, M., Pieper, L., Chew, G.-L. & Schier, A. F. A Massively Parallel Reporter Assay of 3′ UTR Sequences Identifies In Vivo Rules for mRNA Degradation. Mol. Cell 68, 1083–1094.e5 (2017).

56. Doni Jayavelu, N., Jajodia, A., Mishra, A. & Hawkins, R. D. Candidate silencer elements for the human and mouse genomes. Nat. Commun. 11, 1061 (2020).

57. Zhang, F. & Lupski, J. R. Non-coding genetic variants in human disease. Hum. Mol. Genet. 24, R102–R110 (2015).

58. Ward, L. D. & Kellis, M. HaploReg: a resource for exploring chromatin states, conservation, and regulatory motif alterations within sets of genetically linked variants. Nucleic Acids Res. 40, D930–D934 (2012).

59. Untergasser, A. et al. Primer3—new capabilities and interfaces. Nucleic Acids Res. 40, e115–e115 (2012).

60. Grieger, J. C., Choi, V. W. & Samulski, R. J. Production and characterization of adeno-associated viral vectors. Nat. Protoc. 1, 1412–1428 (2006).

61. Gaspar, J. M. NGmerge: merging paired-end reads via novel empirically-derived models of sequencing errors. BMC Bioinformatics 19, 536 (2018).

62. Li, H. Aligning sequence reads, clone sequences and assembly contigs with BWA-MEM. *ArXiv13033997 Q-Bio* (2013).

63. Li, H. et al. The Sequence Alignment/Map format and SAMtools. Bioinformatics 25, 2078– 2079 (2009).

64. Broussard, G. J., Unger, E. K., Liang, R., McGrew, B. P. & Tian, L. Imaging Glutamate with Genetically Encoded Fluorescent Sensors. in Biochemical Approaches for Glutamatergic Neurotransmission (eds. Parrot, S. & Denoroy, L.) 117–153 (Springer, 2018). doi:10.1007/978-1-4939-7228-9_5.

65. Preibisch, S., Saalfeld, S. & Tomancak, P. Globally optimal stitching of tiled 3D microscopic image acquisitions. Bioinformatics 25, 1463–1465 (2009).

66. Schindelin, J. et al. Fiji: an open-source platform for biological-image analysis. Nat. Methods 9, 676–682 (2012).

